# Progesterone induces meiosis through two obligate co-receptors with PLA2 activity

**DOI:** 10.1101/2023.09.09.556646

**Authors:** Nancy Nader, Lama Assaf, Lubna Zarif, Anna Halama, Sharan Yadav, Maya Dib, Nabeel Attarwala, Qiuying Chen, Karsten Suhre, Steven S. Gross, Khaled Machaca

## Abstract

The steroid hormone progesterone (P4) regulates multiple aspects of reproductive and metabolic physiology. Classical P4 signaling operates through nuclear receptors that regulate transcription. In addition, P4 signals through membrane P4 receptors (mPRs) in a rapid nongenomic modality. Despite the established physiological importance of P4 nongenomic signaling, the details of its signal transduction cascade remain elusive. Here, using *Xenopus* oocyte maturation as a well-established physiological readout of nongenomic P4 signaling, we identify the lipid hydrolase ABHD2 (α/β hydrolase domain-containing protein 2) as an essential mPRβ co-receptor to trigger meiosis. We show using functional assays coupled to unbiased and targeted cell-based lipidomics that ABHD2 possesses a phospholipase A2 (PLA2) activity that requires mPRβ. This PLA2 activity bifurcates P4 signaling by inducing clathrin-dependent endocytosis of mPRβ, resulting in the production of lipid messengers that are G-protein coupled receptors agonists. Therefore, P4 drives meiosis by inducing an ABHD2 PLA2 activity that requires both mPRβ and ABHD2 as obligate co-receptors.

**Significance Statement:** Nongenomic progesterone signaling is important for many physiological functions yet the details of its signaling remain elusive. Here we define the early signaling steps downstream of membrane progesterone receptor β (mPRβ) during *Xenopus* oocyte meiosis. We show that progesterone requires two cell membrane receptors to work in unison to signal. The co-receptor complex possesses lipase activity that produces lipid messenger and induces receptor endocytosis to trigger meiosis progression. Our findings have broad physiological implications because nongenomic progesterone signaling operates in many tissues and regulates reproduction and metabolism.

## Introduction

Progesterone (P4) signaling is critical for the regulation of female reproduction, sperm activation, neuronal function, and modulation of the immune system (1–3). The classical mode of P4 signaling is through nuclear receptors (nPRs) that act as transcription factors and modulate gene expression, resulting in cellular responses on a relatively slow time scale (4). In addition, P4 mediates rapid signal transduction that is independent of transcription and has thus been referred to as nongenomic. This signaling modality can be mediated by membrane progesterone receptors (mPRs) that are members of the progestin and AdipoQ receptors family (PAQR), and consist of 11 receptors with 5 being specific to P4: PAQR5 (mPRγ), PAQR6 (mPRδ), PAQR7 (mPRα), PAQR8 (mPRβ) and PAQR9 (mPRɛ) (5). Several lines of evidence support the importance of nongenomic mPR-dependent signaling, including results from studies of nPR knockout mouse lines, the speed of the signal transmission following P4 treatments, and the observed activity of membrane impermeant BSA-coupled P4 (3, 6–8). Currently, mPR-mediated signaling is recognized as an important regulator of many biological functions in the nervous and cardiovascular systems, female and male reproductive tissues, intestine, immune and cancer cells, and glucose homeostasis (3, 6, 9–11). Thus, mPRs are emerging as potential clinical targets for hypertension, reproductive disorders, and neurological diseases (12). However, their downstream signaling events are not well defined.

mPRs are integral membrane proteins with predicted 7-transmembrane domains, a cytoplasmic N-terminus and an extracellular C-terminus, which were first identified in fish ovaries almost 2 decades ago (5, 13, 14). This topology is opposite to that of G-protein coupled receptors (GPCRs), yet there is an abundance of evidence for G protein activation downstream of mPRs, primarily Gαi and also Gβγ (15). However, there are conflicting reports as to whether mPRs signal through trimeric G-proteins or other modalities that remain to be defined (9, 16).

The anuran *Xenopus laevis* is a particularly suitable model to study P4 nongenomic signaling for two reasons: 1) the oocyte is transcriptionally silent, so signaling can only occur through the nongenomic arm; 2) P4 is known to trigger *Xenopus* oocyte maturation through the activation of mPRβ (PAQR8) (14, 17). Fully grown vertebrate oocytes arrest at prophase I of meiosis for prolonged periods before undergoing maturation in preparation for fertilization (18). Importantly, P4 triggers re-entry into meiosis as well as a series of cytoplasmic differentiation steps that allow the egg to become fertilization-competent, able to activate in response to sperm entry, and to support early embryogenesis (19). Signaling downstream of mPRβ ultimately leads to activation of maturation promoting factor (MPF), composed of Cdk1 and cyclin B, that serves as the primary kinase which triggers oocyte entry into M-phase (20). MPF is activated non-reversibly through two signaling cascades, Plk-Cdc25C and the MAPK cascade (Fig. 1A).

**Figure 1.**
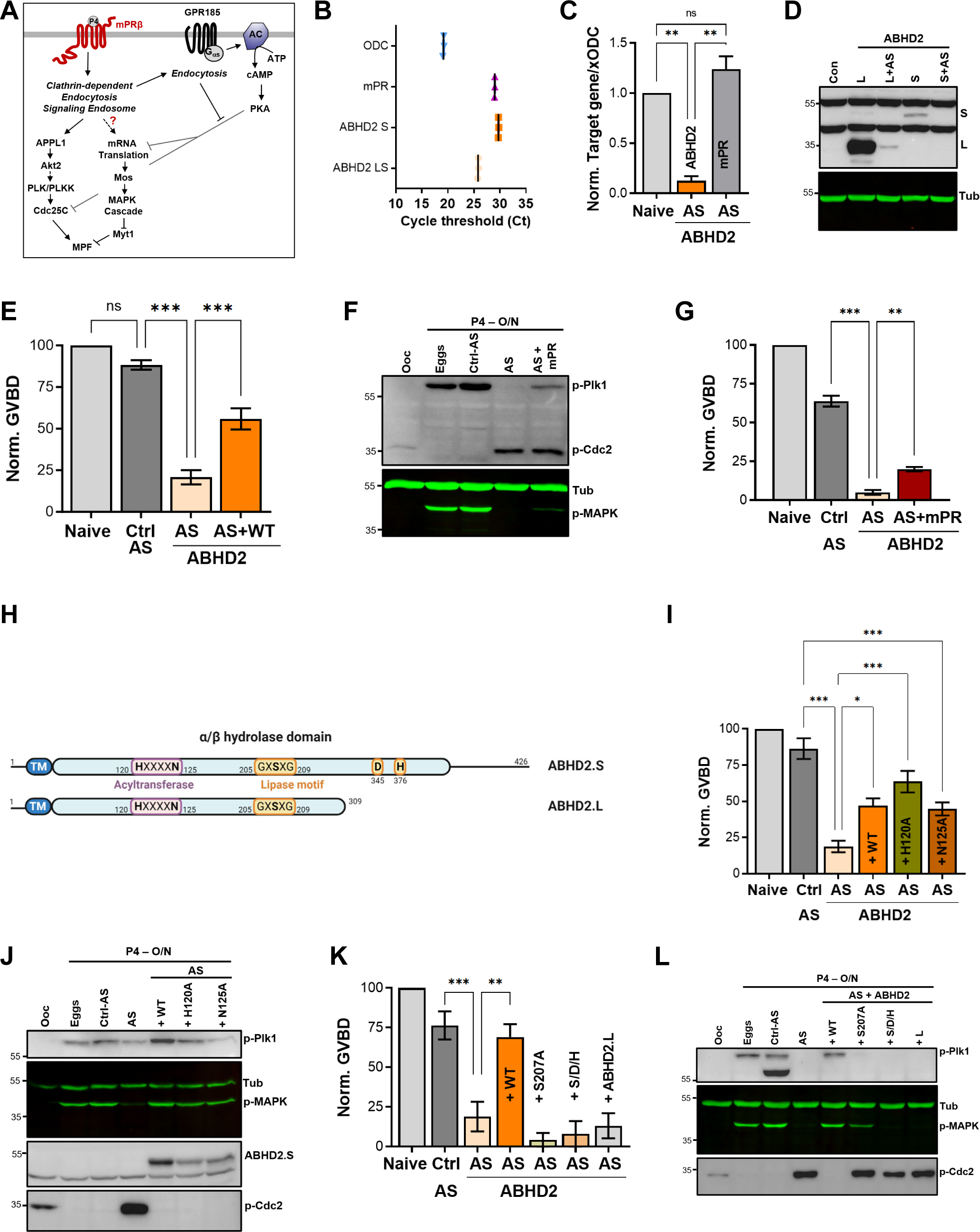
ABHD2 is required for P4-induced oocyte maturation. **A.** Signaling events downstream of mPRβ after progesterone (P4), leading to oocyte maturation. The role of GPR185 is also depicted. **B.** mRNAs transcripts levels of mPRβ, ABHD2.S, ABHD2.LS and *Xenopus* Ornithine decarboxylase (xODC) in oocytes using the Cycle threshold (Ct) generated from real time PCR. **C.** ABHD2 knockdown experiments. Oocytes were injected with specific ABHD2 antisense (AS) oligonucleotides and incubated at 18 °C for 24 hours. RNAs were prepared and analyzed by RT-PCR to determine the efficacy of ABHD2 knockdown as compared to naïve oocytes and mPRβ RNAs levels. Data are expressed as relative RNAs levels of ABHD2 and mPRβ mRNA after normalizing to xODC as a house keeping gene. **D.** Representative WB of ABHD2 in naive and oocytes over-expressing ABHD2.L or ABHD2.S, following ABHD2 antisense (AS) oligonucleotide injection. Tubulin is shown as a loading control. **E.** Oocyte maturation following injection of control antisense (Ctrl AS) or ABHD2 antisense (AS) with or without overexpression of ABHD2.S (AS+WT) and normalized to P4-treated naïve oocytes condition (Naive), (mean ± SEM; n = 7 independent female frogs, ordinary one-way ANOVA). **F.** Representative WB of MAPK, Plk1 and Cdc2 phosphorylation from untreated oocytes, eggs matured by overnight (O/N) treatment with P4, oocytes injected with control antisense (Ctrl AS) or ABHD2 antisense (AS) with or without overexpression of mPRβ. Tubulin is shown as a loading control. **G.** Oocyte maturation in oocytes injected with control antisense (Ctrl AS) or ABHD2 antisense (AS) with or without overexpression of mPRβ and normalized to P4-treated naïve oocytes (mean ± SEM; n = 3 independent female frogs, ordinary one-way ANOVA). **H.** Schematic representation of *Xenopus* ABHD2.S and .L domains, including the acyltransferase and lipase motifs. **I/K.** Oocyte maturation in oocytes injected with control antisense (Ctrl AS) or ABHD2 antisense (AS) with or without overexpression of ABHD2.S wild type (AS+WT) and the different ABHD2.S mutants as indicated, and normalized to P4-treated naïve oocytes (mean ± SEM; n = 3 independent female frogs, ordinary one-way ANOVA). **J/L.** Representative WB of MAPK, Plk1 and Cdc2 phosphorylation from untreated oocytes, eggs matured by overnight (O/N) with P4, oocytes injected with control antisense (Ctrl AS) or ABHD2 antisense (AS) with or without overexpression of the different ABHD2.S mutants as indicated on the panel. Tubulin is shown as a loading control.

In addition, oocyte meiotic arrest is maintained by high levels of cAMP-PKA through the action of a constitutively active GPCR, GPR185 (21, 22). PKA blocks maturation by inhibiting both arms of the pathway: translation and Cdc25C (Fig. 1A) (23, 24). Interestingly, GPR185 is internalized in response to P4, which inhibits its ability to activate adenylate cyclase and thus renders the oocyte more permissive for maturation (22). Several studies have detected a drop in cAMP levels in response to P4 using ELISA (22, 25–28), but these findings could not be replicated using various reporters for cAMP and PKA at the single-cell level in real time (28). Furthermore, the different sensors did not detect a global drop in cAMP in response to P4, and P4-dependent maturation proceeds even when cAMP is high (28). This argues that P4 signals through parallel pathways that can overwhelm the cAMP-PKA-dependent inhibition.

We have previously shown that mPRβ signals through APPL1 and AKT2 to activate Plk and MPF by forming a complex of activators within signaling endosomes (14). Importantly, enrichment of mPRβ within endosomes is sufficient to induce maturation in the absence of P4, arguing that a primary function of P4 is to induce clathrin-dependent endocytosis of mPRβ (14).

The lipase α/β hydrolase domain-containing protein 2 (ABHD2), an integral plasma membrane protein belonging to the ABHD family, was identified as a membrane progesterone receptor in human sperm (12, 29) ABHD2 acts a monoacyl glycerol lipase (MAGL) that hydrolyses 2- arachidonoylglycerol (2-AG) forming arachidonic acid (AA) and glycerol and leads to sperm activation (29). ABHD2 has also been implicated in follicular maturation in mammals (30), vascular smooth muscle migration, and in pulmonary emphysema (31, 32). In addition, ABHD2 was proposed to function as a triacylglycerol (TAG) lipase (33), and shown to localize to the ER where it regulates Ca^2+^ release (34).

Here we show that ABHD2 is required for the P4-induced release of oocyte meiotic arrest. Using untargeted lipidomic analyses, we detect broad downregulation of glycerophospholipid (GPL) and sphingolipid lipid metabolites in association with the enrichment of a key bioactive lipid messengers that include prostaglandins (PGs), lysophosphatidic acid (LPA), and potentially sphingosine-1-phosphate (S1P). Importantly, we show that ABHD2 acts as a PLA2 that requires mPRβ to generate the aforementioned lipid messengers. The ABHD2 evoked PLA2 activity also triggers mPRβ endocytosis, which as we have previously shown is sufficient to initiate oocyte maturation (14). To our knowledge this is the first example of an α/β hydrolase that requires a heterologous co-receptor for activation.

## Results

### ABHD2 is required for P4-induced oocyte maturation

While reviewing the expression levels of various progesterone receptors in the *Xenopus* ovary, we noticed high levels of expression of ABHD2 in the oocyte and egg (Supplemental Fig. 1A). *Xenopus laevis* is tetraploid so it expresses two different versions of each gene, called S and L. There is significant sequence conservation between xABDH2.S and xABDH2.L with the human isoform, expect that xABHD2.L is truncated at position 309 (Supplemental Fig. 1B). Using primers specific to xABDH2.S and ones that amplify both the S and L isoforms, we confirmed that ABHD2 RNA is expressed in the oocyte at levels similar to those of mPRβ RNA (Fig. 1B).

To test whether ABHD2 plays a role in oocyte maturation, we downregulated its expression by injecting anti-ABHD2 (L/S) oligos, which resulted in significant reduction of ABHD2 RNA levels without affecting the levels of mPRβ RNA (Fig. 1C). We could not detect endogenous oocyte ABHD2 using Western blots (WB) most likely due to low expression levels, as we could readily detect overexpressed ABHD2 (Fig. 1D). We thus tested the efficacy of the antisense (AS) approach on overexpressed ABHD2 (either L or S). Figure 1D shows effective reduction of both isoforms (Fig. 1D) within 24 hours after antisense injection, arguing for a short ABHD2 half-life on the order of hours. Interestingly, knocking down ABHD2 expression inhibited oocyte maturation in response to P4 (Fig. 1E), and was coupled to blocking Plk1, MAPK, and MPF activation (Fig. 1F).

This inhibition was significantly reversed by overexpression of ABHD2 (Fig. 1E), confirming that the specific knockdown of ABHD2 – and not off-target effects – mediated AS action. Overexpression of mPRβ in ABHD2 antisense treated oocytes (Supplemental Fig. 1C) was less effective at rescuing oocyte maturation although it did have a significant effect (Fig. 1G), and partially rescued Plk1 and MAPK activation but not MPF (Fig. 1F). These data argue that both mPRβ and ABHD2 are required to sufficiently activate the kinase cascades downstream of P4 to induce oocyte maturation.

### ABHD2 lipase domain is required for P4-induced oocyte maturation

ABHD2 is a serine hydrolase that belongs to the α/β hydrolase family with a conserved lipid hydrolase domain GxSxG with residues D345 and H376 being important for lipase activity based on sequence homology (Fig. 1H). In addition, as with other members of the α/β hydrolase family, *Xenopus* ABHD2 has an acyltransferase domain that is close to the human HxxxD consensus but with Asn replacing the Asp residue (Fig. 1H). As ABHD2 has been shown to function as a P4- dependent hydrolase that is important for sperm activation (29), we wondered whether ABHD2 lipase activity is similarly important for oocyte maturation.

To evaluate this possibility, we replaced the conserved functionally relevant residues in the HxxxD acyltransferase domain motif (i.e., H120 and N125) with Ala and tested the mutants’ ability to induce oocyte maturation in response to P4. Both the H120A and N125A mutants induced oocyte maturation to similar levels as those observed with WT ABHD2 in oocytes injected with ABHD2- AS (Fig. 1I), and this rescue was associated with effective expression of both mutants and activation of Plk1, MAPK, and MPF in eggs (Fig. 1J). These findings demonstrate that the ABHD2 acyltransferase motif is not required for maturation.

To test for the role of ABHD2 lipid hydrolase activity in oocyte maturation, we mutated all three catalytically relevant residues: S207 within the GxSxG motif, D345, and H376 to Ala (S/D/H mutant) or just S207 to Ala. Effective expression of either the S/D/H or S207A mutants (Supplemental Fig. 1D), did not induce maturation in oocytes where endogenous ABHD2 was knocked down (Fig. 1K). Neither mutant activated Plk1 or MPF, but expression of S207A led to low level MAPK activation (Fig. 1L) that was not associated with maturation (Fig. 1K). We further tested whether the shorter ABHD2.L (309 residues) is functional as it is missing the C-terminal end of the α/β hydrolase domain including both D345 and H376. Notably, ABHD2.L did not induce oocyte maturation in response to P4 (Fig. 1K) nor did it activate Plk1, MAPK, or MPF (Fig. 1L). These data show that all three residues involved in lipase activity (S207, D345, H376) are critical for ABHD2 P4-mediated oocyte maturation.

### Lipidomics during oocyte maturation

The requirement for the lipase domain within ABHD2 for P4-dependent oocyte maturation suggests a potential role for lipid messengers downstream of P4 to release oocyte meiotic arrest. There is support for this idea in the literature from several early studies implicating lipid messengers in oocyte maturation, although there has been little consensus regarding specific pathways, lipases, or lipid mediators (35). To globally assess oocyte lipid profiles in response to P4, we performed unbiased mass-spectrometry based lipidomics and metabolomics (using the Metabolon CLP and HD4 platforms) at two time points after P4: 5 min to assess rapid changes in lipid abundances, and 30 min, a ‘point of no return’ where the majority of oocytes in the population commit to maturation (22). Furthermore, to be able to differentiate between pathways that are activated specifically through mPRβ or ABHD2, we performed the profiling on oocytes following mPRβ or ABHD2 knockdown.

The role of lipid signaling in oocyte maturation remains poorly defined, partly due to technical limitations in early studies, but also as a result of the complexity and transient nature of lipid signals (35). An additional confounding factor in oocyte lipidomics is that oocytes in the population mature in an asynchronous fashion in response to P4. This asynchrony makes the identification of transient lipid messengers challenging. Finally, metabolomic profiles depend on environmental factors, basal metabolic rates, diet, and physical activity and are as such quite variable. We observe this variability as metabotypes that are specific to individual frog donors in the principal component analysis in data from both the HD4 and CLP platforms (Supplemental Fig. 2A).

To minimize these technical and biological confounding factors, we performed the metabolomics analysis on groups of 10 oocytes at each time point and looked for relative changes in specific metabolites over time for each donor (Fig. 2A). That is the metabolic profile at T0 served as the baseline for each frog and we looked for changes in each metabolite at the 5- and 30-min time points after P4 addition. Furthermore, we averaged data from specific metabolites at the sub-pathway level to identify changes that are consistent among the 3 donor frogs tested. Data at the specific metabolite levels for both the CLP and HD4 platforms are presented in Supplementary Tables 1-6.

**Figure 2.**
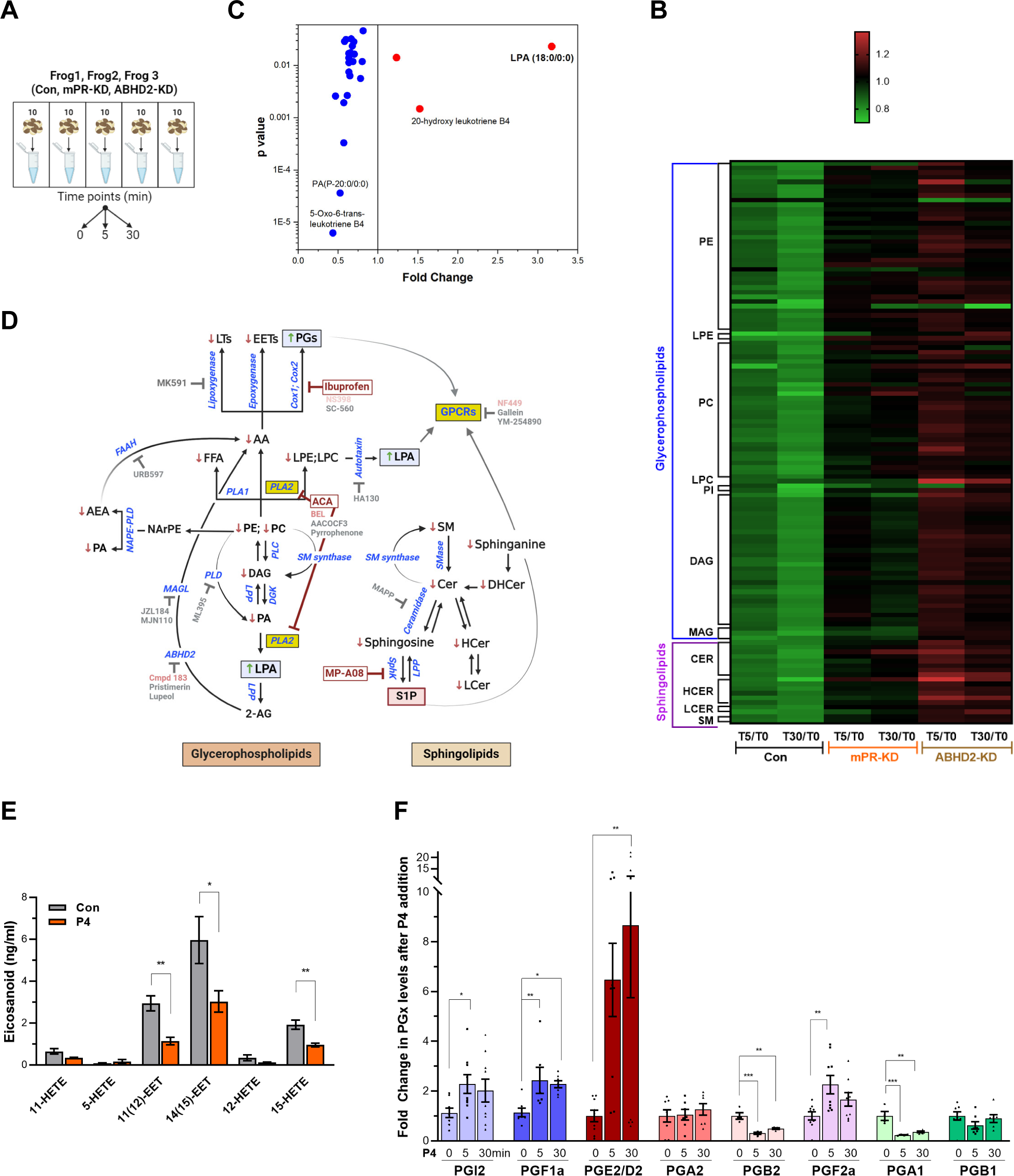
Metabolomics and lipidomics **A. Summary of the experimental setup.** Naïve oocytes (Con) and oocytes injected with either mPR (mPR-KD) or ABHD2 antisense (ABHD2-KD) were treated for 5 or 30 min with P4. For each condition and time point, 5 replicates of 10 pooled oocytes each were collected. The experiment was repeated using 3 donor females. **B.** Heatmap of fold changes for metabolites that were changed significantly (p<0.05) at either the 5 min (T5) or 30 min (T30) time points in response to P4 as compared to untreated oocytes (T5/T0 and T30/T0) for naive (Con), mPR-KD, and ABHD2-KD oocytes. Metabolites are clustered at the levels of glycerophospholipids and sphingolipids and then at the pathway level as follows: PE (Phosphatidylethanolamines), LPE (Lysophosphatidylethanolamines), PC (Phosphatidylcholines), LPC (Lysophosphatidylcholine), PI (Phosphatidylinositols), DAG (Diacylglycerols), MAG (Monoacylglycerols), CER (Ceramides), HCER (Hexosylceramides), LCER (Lactosylceramides), and SM (Sphingomyelins). Supplemental tables 1-4 show the means and p value for each ratio fold change. **C.** Distribution of metabolites that are reduced (blue) or increased (red) significantly in single naïve oocytes 30 min after P4. Fold change and p values were calculated from 10 individual oocytes at each time point. The raw data is listed in Supplemental Table 7. **D.** Summary of the changes in sphingolipids and glycerophospholipids after progesterone treatment. Increase (upward green arrow) or decrease (downward red arrow) in metabolites levels is noted. Tested chemical inhibitors are also shown. Strong inhibitors are indicated in red, weak inhibitors in pink, and drugs that do not inhibit oocyte maturation in gray. Generated using Biorender. **E.** Levels of EETs and HETEs before and after P4 treatment in single oocytes. Naïve oocytes were incubated with either ethanol or P4 10^-5^M for 30 min. 20 single oocytes per condition were collected and used for analysis. 5-oxoETE and 8(9)- EET were detected in 1 or 2 samples respectively, so they were not included in the statistical analyses although both were lower following P4 treatment. **F.** Levels of prostaglandins before (0 min) and after P4 treatment at 5 min and 30 min time points in naïve oocytes. Per each condition, 10 replicates were collected containing 10 pooled oocytes each. The following eicosanoids were not detected in either group: 6kPGF1α, PGF2α, PGE2, TXB2, PGD2, PGA2, PGJ2, 15- deoxyPGJ2, 12-HHTrE, 11-dehydro TXB2, LTB4, LTC4, LTD4, LTE4, 20-hydroxy LTB4, 20- carboxy LTB4, 5(6)-DiHETEs, LXA4, LXB4, 5(6)-EET, 5(6)-DiHET, 8(9)-DiHET, 11(12)-DiHET, 14(15)-DHET, 20-HETE. Similar results were obtained from individual oocyte samples (see Supplemental Table 7). Data are normalized to the PG levels at time zero. Example of the raw abundance of individual PG species is shown in Supplemental Fig. 2C.

Lipidomics analyses on the CLP platform shows downregulation in response to P4 of multiple sphingolipid species, including ceramide, hexosylceramide, lactosylceramide, and sphingomyelin at both the 5 min and 30 min time points (Fig. 2B and Supplemental Tables 1-6). A similar decrease in multiple glycerophospholipid species, including phosphatidylethanolamine (PE), phosphatidylcholine (PC), lysoPE, lysoPC, as well as glycerides including monoacylglycerol (MAG), and diacylglycerol (DAG) is observed (Fig. 2B and Supplemental Tables 1-6). The HD4 platform analysis supports and extends these findings although at a lower level of granularity in terms of specific lipid species showing downregulation of multiple fatty acid species including arachidonic acid (AA)(Supplemental Fig. 2B). Note that most of these changes at the specific biochemical levels are relatively small (20-25% decrease) yet they gain relevance through their combined trend through multiple experiments at the levels of multiple metabolites within the same pathway (Supplemental Tables 1-6).

Importantly, knockdown of either ABHD2 or mPRβ eliminated the downregulation of sphingolipids and glycerophospholipids in response to P4 at both the 5- and 30-min time points (Fig.2B and Supplemental Fig. 2B).As oocytes with mPRβ or ABHD2 knockdown do not mature in response to P4, the lipidomics data argue that the changes observed in the lipid metabolites are important for oocyte maturation; and that both mPRβ and ABHD2 are required to initiate the lipase activities underlying these changes. The fact that many of those changes are observed early on after P4 addition (5 min) argues that they represent the initial signaling step downstream of P4 to commit the oocyte to meiosis.

We were initially puzzled by the global lipidomics profiles as they showed mostly downregulation of both GPL and sphingolipid species. If the cells are indeed in broad lipid catabolism mode, one would expect enrichment in some lipid end products. However, because oocyte maturation is asynchronous, changes in individual lipid species may be masked when several oocytes are pooled as in our lipidomics analyses. To get better insights into changes at the single oocyte level, we developed approaches (see Methods) to extract and analyze metabolites at the single oocyte level at the 30 min post-P4 time point. Single oocytes data showed a 3-fold increase in LPA and replicated the downregulation of GPL and sphingolipid species observed in the pooled analysis (Fig. 2C). The lipidomics changes observed in response to P4 are summarized in Figure 2D along both the GPL and sphingolipid pathways by an arrow up (red) or down (green) next to each detected lipid species.

Some end products of GPL and SL metabolism such as eicosanoids and sphingosine 1 phosphate (S1P) are not well suited to the standard extraction protocols used for our global metabolomics studies. Therefore, to enrich for these end products we performed targeted extraction and MS analyses against validated standards for eicosanoids and S1P, which would be the most likely end metabolites enriched based on the downregulation profile of other metabolites throughout the GPL and SL pathways (Fig. 2D). AA is a precursor for eicosanoids, which are produced through three main enzymatic pathways: 1) cyclooxygenases (cox1/cox2) action results in the formation of prostaglandins; 2) lipoxygenases produce leukotrienes; and 3) cytochrome P450 enzymes, ω- hydrolases and epoxygenases, yield hydroxyeicosatetraenoic acids (HETEs) and epoxyeicosatrienoic acids (EETs), respectively (Fig. 2D).

We could not detect any leukotrienes in the oocyte samples despite testing for multiple species (LTB4, LTC4, LTD4, LTE4, 20-hydroxy LTB4, 20-carboxy LTB4) but could observe a downregulation in their precursor hydroxy-eicosatetraenoic acids (HETE) (Fig. 2E). We did however detect a decrease in LTB4 derivatives in the single oocyte MS analyses (Fig. 2B). We also observed decreases in two epoxyeicosatrienoids (EET) from single oocyte samples (20 oocytes at 30 min post P4) (Fig. 2E). In contrast to leukotrienes and lipoxins, the trend in prostaglandins (PG) was the opposite with increases observed in multiple species, including PGI2, PGF1a, PGE2/PGD2, and PGF2a (Fig. 2F). Figure 2E shows the relative changes of PG species from different donor females. To illustrate the basal abundance of the different PGs species relative to each other, Supplemental Fig. 2C shows the raw abundance of the different PGs relative to each other. Collectively these targeted metabolomics profiles show that AA is preferentially metabolized by cyclooxygenases at the expense of lipoxygenases/epoxygenases to produce PGs in response to P4 (Fig. 2D).

From the metabolomics analyses of the SL pathway, sphingosine-1-phosphate (S1P) is predicted to increase in response to P4, as all other SL metabolites decrease in response to P4 (Fig. 2D). We therefore directly tested S1P levels in the oocyte in response to P4 at 5 and 30 min but did not detect any significant change (Supplemental Fig. 2D). This implies that either S1P does not change in response to P4, or alternatively that the S1P produced in response to P4 diffuses out of the oocyte. This is possible as S1P is membrane permeant and acts on cell surface receptors from the extracellular side (36). We tested this possibility directly below by assessing the role of S1P receptors in oocyte maturation.

### Pharmacological validation of the metabolomics findings

The metabolomics findings from extensive analyses using multiple platforms and extraction procedures to enrich and detect specific lipid species argue for an important role for lipid messengers in inducing oocyte maturation. This is further supported by the fact that when either mPRβ or ABHD2 are knocked down the lipid changes disappear. However, these metabolomics studies are correlative and do not address cause-effect relationships. To test whether activation of specific lipases is required for oocyte maturation, we used pharmacological tools to inhibit specific enzymes predicted to be important for maturation based on our metabolomic analyses (Fig. 2D).

Along the GPL metabolic arm, the lipidomics data show downregulation of upstream metabolites with enrichment of LPA and prostaglandins (PGs) (Fig. 2). This argues for a central role for PLA2 activity as it would produce AA as a precursor for PGs and LPC/LPE as precursors for LPA. To test for a role for PLA2 in maturation, we used a broad-spectrum inhibitor, ACA, as PLA2s represent a large family of enzymes (up to 16 groups) with disparate activation modes, Ca^2+^- dependency, and subcellular localization (secreted or cytosolic) (37, 38). ACA completely inhibited maturation with an IC50 of 3.8x10^-6^ M (Fig. 3A) in line with its reported PLA2 IC50 value (5x10^-6^ M, see Table 1). ACA treatment blocked Plk1, MAPK, and MPF activation explaining the maturation inhibition (Fig. 3B). Given the robust inhibition with ACA, we tested other PLA2 inhibitors, including AACOF3, bromoenol lactone (BEL), and pyrrophenone to get insights into the PLA2 isoform involved. BEL inhibited oocyte maturation with an IC50 of 3.9x10^-5^ M (Fig. 3A), by mainly blocking the Plk1 pathway (Fig. 3C). The BEL IC50 was about 5-fold higher than its reported inhibitory potency of 8x10^-6^ M (Table 1). The other two PLA2 inhibitors were ineffective (Fig 3A). It is difficult to pinpoint a particular PLA2 isoform based on the isoform specific profile of these inhibitors (see https://www.sigmaaldrich.com/QA/en/technical-documents/technical-article/protein-biology/protein-expression/phospholipase-a2), arguing that the oocyte’s PLA2 activity required for maturation does not match known PLA2 isoforms. Nonetheless, the inhibitors data support a central role for PLA2 in inducing oocyte maturation downstream of P4. In line with this, pre-treatment of oocytes with mastoparan, a peptide toxin from wasp venom known to activate PLA2, enhanced oocyte maturation by ∼50% in response to low P4 (Fig. 3D).

**Figure 3.**
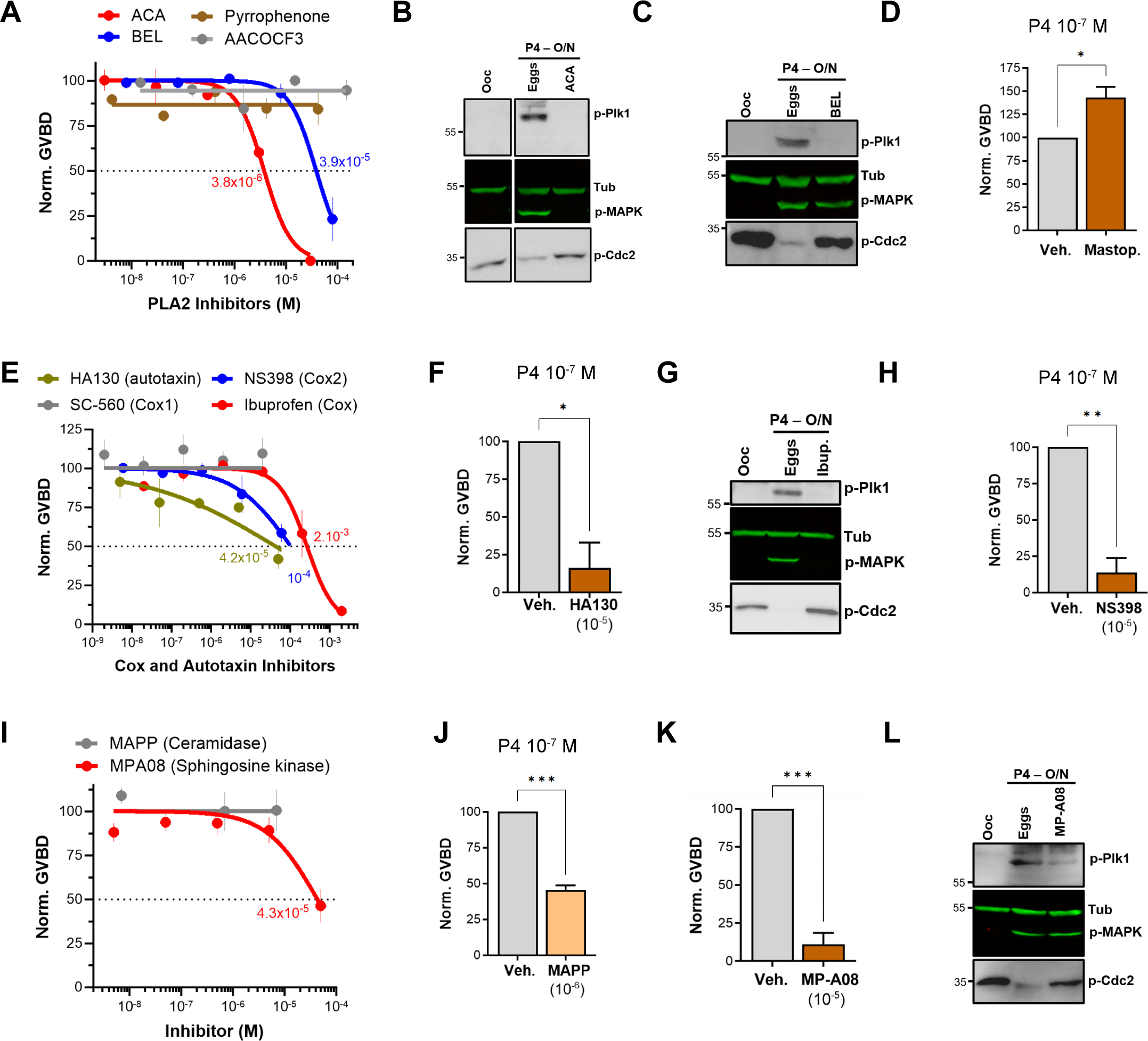
Pharmacological validation of the metabolomics findings. A/E/I. Dose response of inhibition of oocyte maturation for the drugs tested. Oocytes were pre-treated for 2 hours with vehicle or with increasing concentrations of the indicated drugs, followed by overnight treatment with P4 at 10^-5^M. IC50 was calculated using a nonlinear regression fit (mean ± SEM; n = 3 independent female frogs). **D/F/H/J/K.** Drug effect on oocytes maturation at low P4 concentration. Oocytes were pre-treated for 2 hours with the vehicle or with the highest drug concentration from the dose response, followed by overnight treatment with P4 at 10^-7^M. Oocyte maturation was normalized to control oocytes (treated with vehicle) (mean ± SEM; n = 3 independent female frogs for each chemical compound experiment, unpaired t-test). **B/C/G/L.** Representative WB of MAPK, Plk1 and Cdc2 phosphorylation from untreated oocytes, oocytes pretreated with vehicle or the indicated drug for 2 hours then matured overnight (O/N) with P4 (eggs). Tubulin is shown as a loading control.

**Table 1.** Pharmacological inhibitors potency.

| Table 1. Pharmacological inhibitors potency |  |  |  |  |
| --- | --- | --- | --- | --- |
| Inhibitor | Target | IC<sub>50</sub> ± SEM<br>Oocyte<br>Maturation (M) | IC<sub>50</sub> (M)<br>Representative<br>Study | Ratio<br>(Ooc/Lit) |
| ACA | PLA2 | 3.8E-6 ± 5.8E-7 | 5E-6 (54) | 0.76 |
| BEL | PLA2 | 3.9E-5 ± 8E-6 | 8E-6 (55) | 4.9 |
| MP-A08 | Sphingosine Kinase 1 | 4.3E-5 ± 1E-5 | 2.7E-5 (56) | 1.6 |
| HA130 | Autotaxin | 4.2E-5 ± 4.8E-5 | 2.8E-8 (57) | 1,522 |
| Ibuprofen | Cox-2 | 2.7E-4 ± 6E-5 | 3.7E-4 (58) | 0.73 |
| NS398 | Cox-2 | 1E-4 ± 5E-5 | 3.8E-6 (59) | 26.3 |
| NF-449 | GαS | 5.2E-5 ± 3E-5 | 7.9E-6 (60) | 6.5 |

AA could still be produced through the action of fatty acid amide hydrolase (FAAH) independently of PLA2 activation, and PLD activation produces phosphatidic acid (PA) which is a precursor of LPA (Fig. 2F). Inhibition of either FAAH or PLD had no effect on oocyte maturation (Supplemental Fig. 3A), arguing that the precursors for LPA and PGs are produced primarily through PLA2 activation in response to P4 in the oocyte.

Given the inhibitory effect of ABHD2 knockdown on oocyte maturation, we tested the effect of two plant triterpenoids known to inhibit ABHD2 and MAGLs, pristimerin and lupeol (39), as well as MAGL chemical inhibitors JZL184 and MJN110. None of these inhibitors had a significant inhibitory effect on oocyte maturation at high P4 (Supplemental Fig. 3B). However, because pristimerin showed a trend toward inhibition at the highest concentration, we tested its effect at low P4 concentration where it inhibited oocyte maturation but required a concentration of 10^-5^ M (Supplemental Fig. 3C), which is two orders of magnitude higher than its documented IC50 in other systems (39). These results argue against an important role for a monoacylglycerol enzymatic activity, whether ABHD2-dependent or not, in inducing oocyte maturation. We further tested a reported specific ABHD2 inhibitor, compound 183 (10^-4^M), which inhibited oocyte maturation at low and high P4 concentrations (Supplemental Fig. 3D), arguing for a role for ABHD2 in oocyte maturation.

We next focused on enzymes that generate PGs and LPA as the enriched lipid metabolites in response to P4 (Fig. 2D). LPA can be produced through the action of PLA2 on phosphatidic acid (PA) or through autotaxin metabolism of LPE/LPC (Fig. 2D). Blocking autotaxin activity with HA130 repressed oocyte maturation with an IC50 of 4.2x10^-5^ M in the presence of high P4 (Fig. 3E). We therefore tested the effects of HA130 at low P4, where it significantly inhibited maturation by ∼80% (Fig. 3F). However, this inhibition of maturation required high concentrations of HA130 -at least 3 orders of magnitude higher that the reported HA130 IC50, arguing against a major role for the autotaxin pathway in generating LPA in response to P4, and rather supporting LPA production primarily through PLA2 activity.

We then tested the effect of inhibiting cyclooxygenases responsible for PGs production on oocyte maturation. Inhibition of Cox enzymes with ibuprofen effectively blocked maturation with an IC50 of 2.7x10^-4^ M consistent with its IC50 against Cox2 (3.7x10^-4^ M) (Fig. 3E, Table 1). As expected from its effect on maturation, ibuprofen treatment blocked both Plk1 and MAPK activation (Fig. 3G). Specific inhibition of Cox2 using NS398 blocked maturation with an IC50 of 10^-4^ M (Fig. 3E), which is higher than its documented IC50 against Cox2 (3.8x10^-6^ M) (Table 1). Consistently, NS398 at high concentration had a more pronounced inhibitory effect on oocyte maturation (∼85%) at limiting P4 concentrations (Fig. 3H). The Cox1 specific inhibitor SC-560 had no effect on oocyte maturation (Fig. 3E). Collectively these data argue for a role for Cox2 in supporting maturation but not for Cox1.

Interestingly, inhibition of lipoxygenases using MK591 potentiated maturation by ∼20% in the presence of high P4 (Supplemental Fig. 3E), and by 100% at low P4 (Supplemental Fig. 3F). Leukotrienes are decreased in response to P4 (Fig. 2), so presumably inhibition of lipoxygenases diverts AA metabolism toward cyclooxygenase hydrolysis thus increasing PGs production, which would explain the observed effect on oocyte maturation.

### S1P signaling supports oocyte maturation

The lipidomics data show downregulation of most metabolites along the sphingolipid arm of the metabolic pathway with a presumed enrichment of S1P (Fig. 2D). We were however unable to detect an increase of S1P in the oocyte in response to P4 even when using a targeted mass spectrometry-based assay for S1P (Supplemental Fig. 2D). This may be because S1P is readily secreted and signals extracellularly. However, activation of ceramidases could be involved in the sphingolipid changes. mPRs share sequence and structural homology with adiponectin receptors, which have been shown to possess ceramidase activity that requires Zn in the active site that acts on the ceramide amide bond (40). We have previously shown that an mPRβ mutant with all 4 zinc coordinating residues mutated to Ala to abrogate Zn binding (H129,D146,H281,H285A) is functional in inducing oocyte maturation (14). This argues that mPRβ does not require ceramidase activity to release oocyte meiotic arrest. To confirm this finding, we mutated Ser125 in the conserved SxxxH motif in mPRβ, which matches the ceramidase domain in PAQR and AdipoQ receptors (41, 42). The mPRβ-S125A mutant expresses and traffics normally to the cell membrane to similar levels as WT mPRβ (Supplemental Fig. 4A). Furthermore, it signals effectively downstream of progesterone as its overexpression (Supplemental Fig. 4B) rescues mPRβ knockdown with similar efficiency as WT mPRβ (Supplemental Fig. 4C), and activates Plk1, MAPK, and MPF to induce oocyte maturation (Supplemental Fig. 4D). Collectively these results argue against a role for mPRβ-dependent ceramidase activity in inducing oocyte maturation. Hence, changes in sphingolipids might be a consequence of the alterations in other lipid metabolic pathways especially that the metabolism of GPLs is linked to that of sphingolipids through the activity of SM synthase.

If indeed S1P is the end product of sphingolipid metabolism in response to P4, then one would predict activation of sphingosine kinases and concurrently ceramidases to support the metabolic flux. We tested this hypothesis using broad spectrum sphingosine kinase and ceramidase inhibitors. Inhibition of ceramidase activity using D-e-MAPP had little effect on oocyte maturation at high P4 (10^-5^M) (Fig. 3I). In contrast, at limiting P4 (10^-7^ M) MAPP inhibited oocyte maturation by ∼55% compared to untreated oocytes (Fig. 3J). Note that at low P4 (10^-7^ M) oocyte maturation is not complete as only the most primed oocytes respond, and this response varies from frog to frog (10-50% of the maturation observed with high P4). Inhibition of sphingosine kinases using MP-A08 inhibited oocyte maturation with an IC50 of 4.3x10^-5^ Mat high P4 (Fig. 3I and Table 1). At low P4, MP-A08 blocked maturation by 87.5% (Fig. 3K), supporting a role for S1P in oocyte maturation. MP-A08 treatment primarily blocks Plk1 activation and to a lesser extent MAPK (Fig. 3L).

To assess the role of S1P signaling in releasing oocyte meiotic arrest, we targeted S1P receptor (S1PR) isoforms based on their expression in the ovary. Of the 5 S1PR isoforms, only S1PR2 and S1PR3 are expressed in the ovary with higher expression of S1PR3 (43). We first confirmed the expression profile in oocytes using RT-PCR (Fig. 4A). S1PR3 expression levels were higher than S1PR2 (Fig. 4A). We confirmed the efficiency of the S1PR2 primers in the spleen where S1PR2 was shown to be highly expressed (43). We then focused on S1PR3 as the primary S1PR in the oocyte and knocked down its expression using antisense oligos. We validated S1PR3 knockdown at the RNA (Fig. 4B) and protein (Fig. 4C) levels. S1PR3 knockdown blocked oocyte maturation induced by either P4 or the mPR specific agonist OD-02 by ∼50% (Fig. 4D).

**Figure 4.**
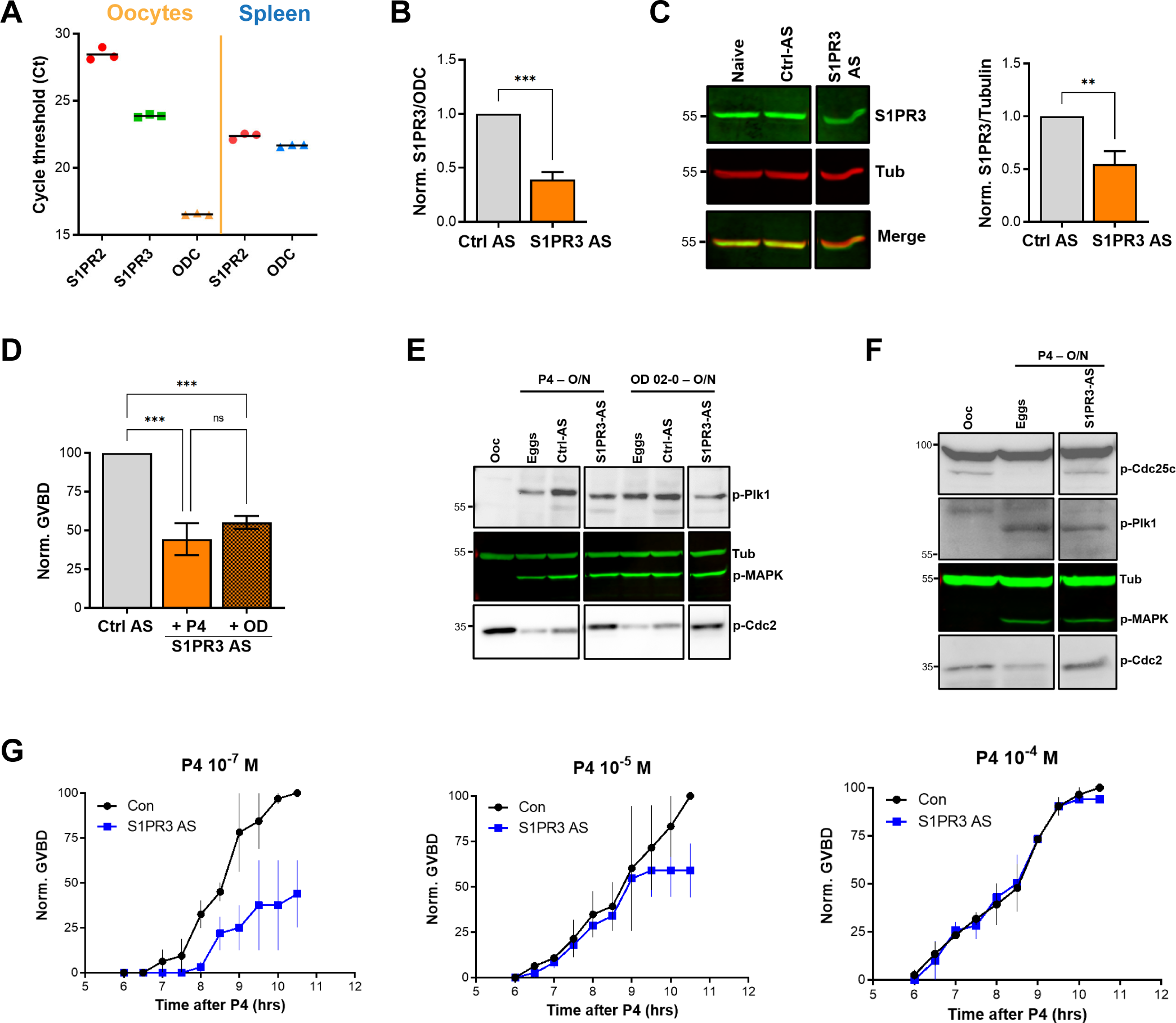
S1P signaling and oocyte maturation. **A.** mRNAs levels of S1PR2, S1PR3, and the house keeping gene Ornithine decarboxylase (ODC) in oocytes and spleen measured as the Cycle threshold (Ct) from real time PCR. **B-C.** S1PR3 knockdown. Oocytes were injected with control antisense (Ctrl AS) or specific S1PR3 antisense oligonucleotides and incubated at 18 °C for 24 hours. RNAs and proteins extracts were prepared and analyzed by RT-PCR and WB to determine the efficacy of S1PR3 knockdown as compared to control oocytes (Ctrl AS). B. Histogram showing the relative RNAs levels of S1PR3 mRNA to xODC. C. Representative WB (left panel) and normalized quantification (right panel) comparing S1PR3 protein levels between naïve, Ctrl AS and S1PR3 AS injected oocytes. Tubulin is shown as a loading control (mean ± SEM; n = 6 independent female frogs). **D.** Oocyte maturation following injection of S1PR3 antisense, normalized to P4 or Org OD 02-0 (OD)-treated oocytes injected with control antisense (Ctrl AS) (mean ± SEM; n = 7 independent female frogs, ordinary one-way ANOVA). **E/F.** Representative WBs of MAPK, Plk1 and Cdc2 (E), as well as CDC25C (F) phosphorylation from untreated oocytes, P4 or OD matured eggs (O/N)D, oocytes injected with control antisense (Ctrl AS) or S1PR3 antisense (S1PR3 AS) and treated O/N with P4 or OD. Tubulin is shown as a loading control. **G.** GVBD-time course after treatment with P4 at the indicated concentrations in oocytes injected with water (Con) or with S1PR3 antisense (S1PR3 AS) (mean ± SEM; n = 2 independent female frogs).

Interestingly, S1PR3 knockdown did not affect P4-mediated activation of MAPK and Plk1 (Fig. 4E), yet P4 was unable to dephosphorylate and activate MPF (Fig. 4E). Plk1 induces maturation by activating the dual-specificity phosphatase Cdc25C, which dephosphorylates Cdc2 to activate MPF and is considered a rate limiting step in MPF activation (44). We therefore tested the effects of S1PR3 knockdown on Cdc25C activation and show that despite the induction of both Plk1 and MAPK, Cdc25C was not activated in response to P4 in S1PR3 knockdown cells (Fig. 4F). We followed Cdc25C activation by assessing its dephosphorylation at Ser287 (Ser216 in human), which is required for its activation (Fig. 4F) (44). The partial oocyte maturation block in S1PR3 KD oocytes argues for a modulatory role for S1P signaling. We tested this hypothesis by assessing the requirement for S1PR3 at increasing P4 dosages that would incrementally induce the multiple arms of the maturation signaling pathways. S1P3R knockdown oocytes matured with significantly less efficiency and with a slower time course at low P4 (10^-7^ M) (Fig. 4G), but interestingly this effect was gradually lost with increased P4 concentrations (10^-5^ and 10^-4^ M) (Fig. 4G). These data argue that the S1P/S1PR3 pathway modulates oocyte maturation, but that by itself it is not essential for maturation as it can be bypassed when other pathways are hyper activated at high P4 concentrations. This is consistent with the production of multiple lipid messengers to induce maturation through several GPCR pathways.

### Role of G-protein coupled receptors (GPCRs) in mediating lipid messenger action

All the enriched lipid metabolites detected from our lipidomics analyses - PGs, LPA, and possibly S1P - act through G-protein coupled receptors (GPCRs). This raises the intriguing possibility that P4 expands its signaling through the production of lipid messengers that act on several GPCRs to mediate the multiple cellular changes that need to occur concurrently during oocyte maturation. This is an attractive possibility as GPCRs have been implicated in *Xenopus* oocyte maturation for decades without conclusive evidence for their involvement, and they have been shown to mediate mPR nongenomic signaling. Therefore, we tested the effects of broad inhibitors of trimeric G-proteins. Blocking Gαs using NF-449 partially inhibited maturation (IC50 5x10^-5^ M) (Supplemental Fig. 4E), whereas blocking either Gβγ or Gαq/11 with gallein and YM-254890 respectively was inefficient at blocking oocyte maturation (Supplemental Fig. 4E). Interestingly, inhibiting Gαs with NF449 primarily blocked Plk1 activation to inhibit MPF and oocyte maturation, and to a lesser extent the MAPK pathway (Supplemental Fig. 4F). At low P4 concentration the inhibitory effect of NF-449 was still observed as well as inhibition by gallein but not YM-254890 (Supplemental Fig. 4G). This argues for a supporting role for Gβγ signaling as well in oocyte maturation. These results support a role for Gαs signaling and potentially Gβγ downstream of P4 through lipid intermediates that require PLA2, Cox2, and SphK activities.

### ABHD2 is a PLA2 that requires mPRβ as a co-receptor

Both the metabolomics and pharmacological data strongly implicate PLA2 activation downstream of P4 to induce maturation. In addition, several lines of evidence argue that this PLA2 activity requires ABHD2: 1) Induction of the signaling cascades downstream of P4 requires both ABHD2 and PLA2; 2) the lipidomics changes downstream of P4 are dependent on both activities; 3) mutations in the ABHD2 lipase domain abolish its ability to induce maturation; 4) ABHD2 has been characterized as a lipase acting on either MAG or TAG (29, 33); and 5) an ABHD2 specific inhibitor (compound 183) inhibits maturation. Collectively, these findings suggest that ABHD2 is associated with a PLA2 activity that is induced in response to P4 to release oocyte meiotic arrest.

To test this hypothesis, we translated ABHD2 and mPRβ *in vitro* in rabbit reticulocyte lysates and assessed whether they possess PLA2 activity. Expression of ABHD2 or mPRβ alone was not associated with any increase in PLA2 activity whether in the presence or absence of P4 (Fig. 5A). Interestingly though, co-expression of both mPRβ and ABHD2 produced a PLA2 activity, but only in the presence of P4 (Fig. 5A). This despite our inability to detect either mPR or ABHD2 expression by Western blots from the reticulocyte lysates reactions. These results argue that both mPRβ and ABHD2 are required for the P4-dependent PLA2 activity, but they do not address which receptor is the catalytic subunit. However, our functional experiment with the ABHD2 hydrolase domain mutants (S/D/H and S207) strongly argues that ABHD2 is the PLA2 catalytic subunit.

**Figure 5.**
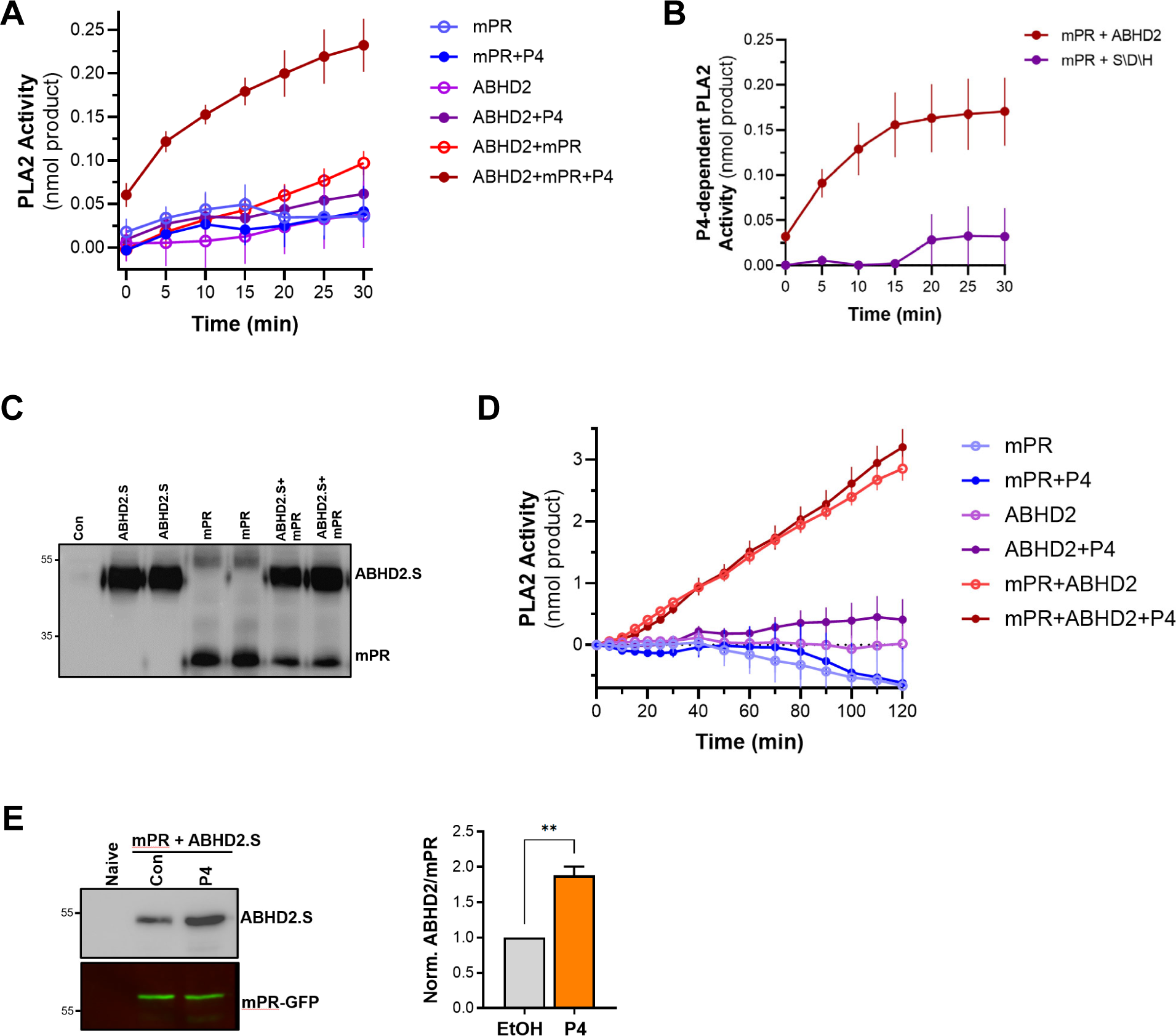
PLA2 activity of the mPRβ-ABHD2 co-receptor complex. **A.** Time-dependent PLA2 activity from reticulocyte lysates expressing mPR, ABHD2.S, or ABHD2.S+mPR in the presence of ethanol as vehicle or P4 10^-5^M (mean ± SEM; n = 3). **B.** Time course of P4-dependent PLA2 activity in reticulocyte lysates expressing mPRβ with either wild-type ABHD2 (ABHD2) or the ABHD2 S207A/D345A/H376A mutant (S/D/H). P4-dependent PLA2 is plotted as the difference in activity in the presence and absence of P4 (Mean ± SEM; n = 3). **C.** Example WB probed with anti-His antibody from tobacco lysates (ALiCE®) alone (Con) and lysates expressing mPRβ or ABHD2 alone or both proteins in duplicated as indicated. **D.** Time-dependent PLA2 activity from ALiCE® lysates overexpressing mPR, ABHD2.S, or ABHD2.S+mPR in the presence of the vehicle ethanol or P4 10^-5^M (mean ± SEM; n = 3). Data are plotted as the rate of production of the lysothiophospholipid product from the beginning of the experiment (0 min) with the rate of the lysates alone subtracted. **E.** Representative immunoprecipitation (IP) WB and quantification of mPR-GFP from oocyte lysates from un-injected oocytes (Naive) or oocytes over-expressing mPR-GFP and ABHD2.S treated for 40 min with Ethanol (EtOH) or P4. *Left panel,* the representative WB membrane is probed for ABHD2 and GFP (mean ± SEM; n = 3 independent female frogs, unpaired t-test).

To directly confirm this prediction, we co-expressed mPRβ with either ABHD2 WT or the S/D/H mutant in reticulocyte lysates and tested their PLA2 activity. Mutating the ABHD2 lipid hydrolase domain (S/D/H mutant) resulted in loss of PLA2 activity when the protein was expressed with mPRβ in the presence of P4 (Fig. 5B). Taken together, these results argue that ABHD2 has PLA2 catalytic activity but only in the presence of mPRβ.

To cross validate our findings from the reticulocyte experiments, we tested in vitro and cell-based expression systems to increase protein yield. We settled on the coupled in vitro transcription/translation tobacco lysates system (ALiCE®) that produces high protein yields and targets expressed proteins to microsomes directly using a melittin signal peptide. This is an advantage for our proteins of interest as they are both integral membrane proteins. The tobacco lysates expression system produced significantly higher yields of both mPR and ABHD2 expression where both proteins were readily detected on Western blots (Fig. 5C). Similar to the reticulocyte system, tobacco lysates expressing ABHD2 or mPRβ alone were not associated increased PLA2 activity whether in the presence or absence of P4 (Fig. 5D). Of note tobacco lysates have endogenous PLA2 activity, which was inhibited by expression of mPRβ alone (Fig. 5D). The PLA2 activity data from tobacco lysates are reported as the rate of increase over time (Fig. 5D). Co-expression of mPRβ and ABHD2 in the tobacco lysates produced significant PLA2 activity that was not only higher than that detected from the reticulocyte lysates but also stable over extended periods of time allowing recordings for over 2 hours (Fig. 5D). Interestingly in this case however the PLA2 activity of the co-receptor complex was independent of P4 (Fig. 5D). Both the reticulocyte and tobacco expression system show that mPRβ is a necessary partner for ABHD2 to mediate its PLA2 activities. However, the reticulocyte co-expression results argue that mPRβ and ABHD2 need P4 to assemble and form a complex to be able to mediate the ABHD2 PLA2 activity. In contrast, when the two proteins are expressed at a high levels and more important targeted specifically to the same microsomal compartment their PLA2 activity becomes P4 independent. We were thus interested in testing whether the two receptors interact in vivo. We expressed mPRβ-GFP in oocytes and tested whether it forms a complex with ABHD2. Immunoprecipitation of mPRβ-GFP pulls down ABHD2 at rest (Fig. 5E), and importantly this interaction is significantly enhanced (1.9±0.2-fold, p=0.002) following P4 treatment (Fig. 5E). This argues that P4 enhances the assembly of the co-receptor complex. Furthermore, co-expression and pull-down experiments show that the non-functional ABHD2 mutants: S/D/H and ABHD2.L interact with mPRβ (Supplemental Fig. 4H), indicating that the interaction between ABHD2 and mPRβ does not require a functional lipase domain.

### ABHD2 and PLA2 activity are required for mPRβ endocytosis and signaling

We have previously shown that mPRβ induces oocyte maturation at the level of the signaling endosome following its clathrin-dependent endocytosis (14). Importantly, mPRβ endocytosis is sufficient to induce maturation in the absence of P4. Since changes in plasma membrane lipid composition are known to modulate endocytosis and membrane components activities, we suspected a role for ABHD2/PLA2 in mPRβ endocytosis in response to P4. Therefore, we quantified mPRβ endocytosis following ABHD2 knockdown. P4 leads to enrichment of mPRβ positive intracellular vesicles, and this enrichment requires ABHD2 as it was lost in ABHD2 knockdown oocytes (Fig. 6A). PLA2 activity was also critical for P4 induced mPRβ endocytosis, which was completely blocked in ACA treated oocytes (Fig. 6B). Interestingly, ACA also inhibited basal endocytosis of mPRβ in the absence of P4, which is apparent as lower vesicular mPRβ in oocytes treated with ACA (Fig. 6B, no P4). This argues that both ABHD2 and PLA2 are required for mPRβ endocytosis and by extension its signaling in response to P4.

**Figure 6.**
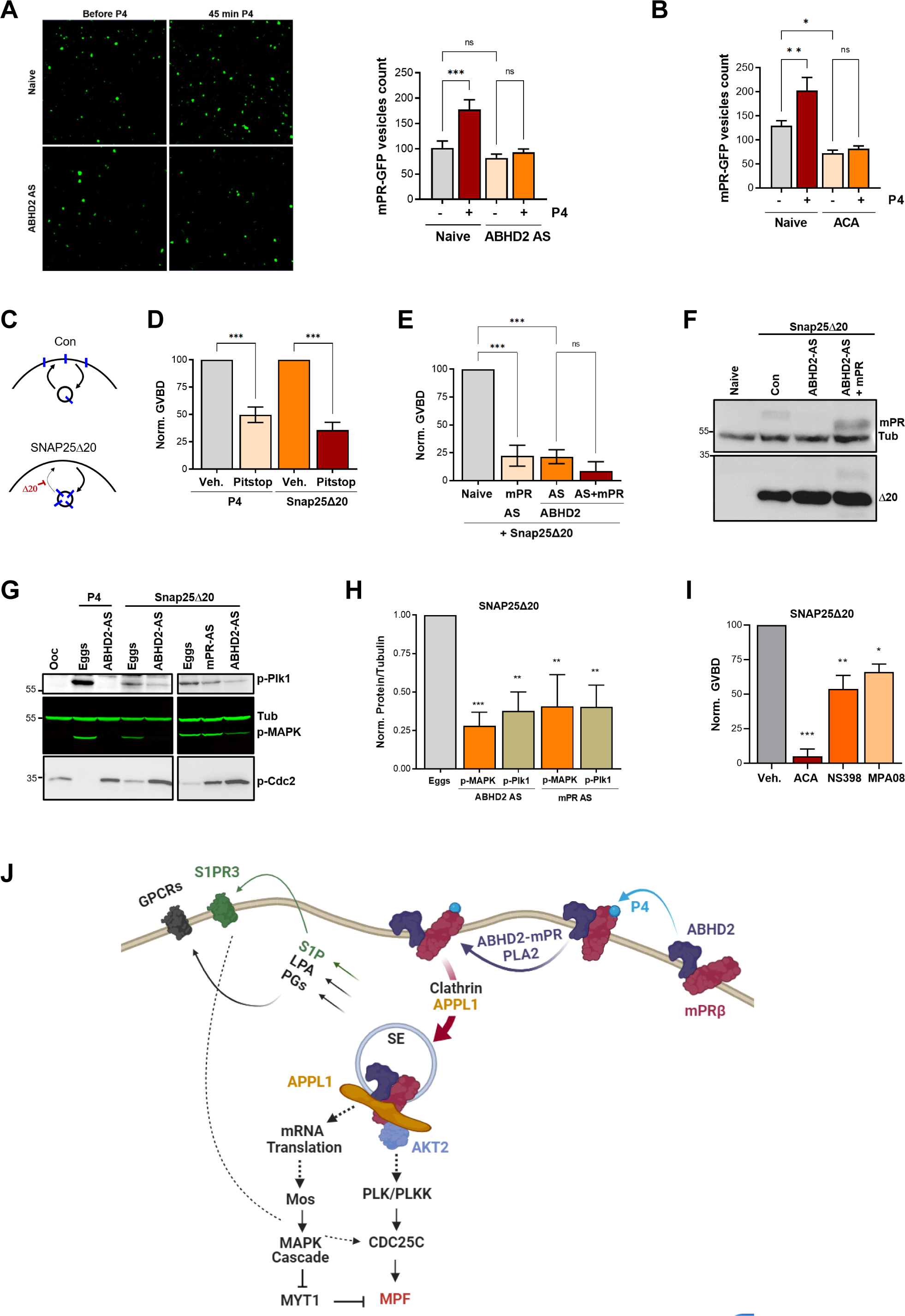
ABHD2 and PLA2 activity are required for mPRβ endocytosis and signaling. **A.** Effect of ABHD2 knockdown on mPR-P4 mediated endocytosis. Oocytes were injected with mPR-GFP RNA (Naïve) and 48 hours later either left untreated or injected with ABHD2 antisense (ABHD2 AS). The following day oocytes were imaged, before and 45 min after P4 treatment*. Left panel,* representative confocal image of mPR-GFP positive vesicles in naïve and ABHD2 AS oocytes, before and after P4. *Right panel,* histogram of mPR-GFP positive vesicles count before and after P4 (mean ± SEM; n = 20 to 21 oocytes per condition, from 3 independent female frogs, ordinary one-way ANOVA). **B.** PLA2 inhibition blocks P4 mediated endocytosis. Vesicle count from oocytes expressing mPR-GFP for 72 hours were pre-treated with vehicle (naïve) or ACA for 2 hours, followed by imaging, before and 45 min after P4 treatment (mean ± SEM; n = 16 to 18 oocytes per condition, from 2 independent female frogs, ordinary one-way ANOVA). **C.** Cartoon depicting the role of SNAP25Δ20 in blocking exocytosis. **D.** SNAP25Δ20-induced oocyte maturation requires clathrin-dependent endocytosis. Naïve oocytes were pre-treated with vehicle or Pitstop (10^-5^ M), followed by overnight treatment with P4 or SNAP25Δ20-mRNA injection. Oocyte maturation in P4 or SNAP25Δ20 injected oocytes normalized to the vehicle condition (mean ± SEM; n = 4 independent female frogs, ordinary one-way ANOVA). **E.** SNAP25Δ20- induced maturation requires ABHD2. Oocytes were injected with mPR antisense (mPR AS) or ABHD2 antisense (AS) with or without mPRβ mRNA (AS+mPR). 48 hours later, oocytes were injected with mRNA to overexpress SNAP25Δ20. The following day, oocyte maturation was measured in mPR AS, ABHD2 AS and ABHD2 AS+mPR oocytes normalized to naive oocytes injected with SNAP25Δ20 (mean ± SEM; n = 3 independent female frogs, ordinary one-way ANOVA). **F.** Representative WB of mPR and SNAP25Δ20 proteins expression in naïve, ABHD2 AS and ABHD2 AS+mPR oocytes. Tubulin is shown as a loading control. **G.** Representative WB of MAPK, Plk1 and Cdc2 phosphorylation from untreated oocytes, P4 matured eggs, or SNAP25Δ20 mRNA injection, and oocytes injected with mPR (mPR AS) or ABHD2 antisense (ABHD2 AS) and treated O/N with P4 or SNAP25Δ20 mRNA injection. Tubulin is shown as a loading control. **H.** Quantification of p-Plk as the ratio of p-PLK/Tubulin and p-MAPK as the ratio of p-MAPK/Tubulin normalized to the ratios in naive eggs (mean ± SEM; n = 4 independent female frogs, ordinary one-way ANOVA). **I.** Oocyte maturation in oocytes pretreated for 2 hours with ACA, NS398 and MP-A08 followed by SNAP25Δ20-mRNA injection and normalized to GVBD in oocytes pre-treated with vehicle followed by SNAP25Δ20-mRNA injection (Veh.) (mean ± SEM; n = 3 independent female frogs, ordinary one-way ANOVA). **J.** Model of the signaling cascade triggered in response to P4 (generated using Biorender).

We were therefore interested in assessing the subcellular distribution of ABHD2 during oocyte maturation to determine whether it is also internalized. We thus tagged it at the N- or C-terminus with mCherry. Unfortunately, both mCh-tagged ABHD2s were not functional as they did not rescue ABHD2 knockdown (Supplemental Fig. 4I) despite efficient expression and binding to mPR (Supplemental Fig. 4H). This prevented us from imaging the trafficking and distribution of ABHD2 during oocyte maturation.

The requirement for ABHD2 to mediate P4-dependent mPRβ endocytosis (Fig. 6A) suggests that this may be the primary function of ABHD2 in response to P4. Under that scenario, P4 activates ABHD2 enzymatic activity which induces mPRβ endocytosis leading to oocyte maturation. To test this hypothesis, we were interested in inducing mPRβ endocytosis in the absence of P4. We have previously shown that a dominant negative SNAP25 mutant missing the last 20 residues (SNAP25Δ20), effectively blocks exocytosis in the oocyte (Fig. 6C); leading to mPRβ enrichment intracellularly and inducing maturation in the absence of P4 (14, 45). Similar to P4, SNAP25Δ20- induced oocyte maturation requires clathrin-dependent endocytosis as it was blocked by clathrin blocker pitstop (Fig. 6D). As expected, SNAP25Δ20-induced oocyte maturation requires mPRβ as it was inhibited in oocytes where mPRβ was knocked down (Fig. 6E). Surprisingly, SNAP25Δ20- induced maturation also requires ABHD2, since oocytes where ABHD2 is knocked down were unable to mature in response to SNAP25Δ20 (Fig. 6E), despite high expression of SNAP25Δ20 (Fig. 6F). We further tested whether mPRβ overexpression allows ABHD2 KD oocytes to mature in response to SNAP25Δ20, but this was not the case (Fig. 6E). This argues that even in the presence of excess mPRβ, its forced internalization using SNAP25Δ20 (Fig. 6C) in the absence of ABHD2 is not sufficient to induce maturation. Therefore, ABHD2 activity is not only required for mPRβ internalization but it is also required at the level of the signaling endosome following mPRβ endocytosis to induce maturation.

Like P4, SNAP25Δ20 activates Plk1, MAPK, and MPF (Fig. 6G) and this induction was significantly inhibited following knockdown of either ABHD2 or mPRβ (Fig. 6G and 6H), showing that both proteins signal through the canonical kinase cascades to induce maturation following internalization even in the absence of P4. Furthermore, in addition to the PLA2 inhibitor ACA, inhibition of Cox2, or sphingosine kinase partially blocked SNAP25Δ20 induced maturation with ACA being the most effective (Fig. 6I), similar to what we observe with P4. ACA blocks basal mPRβ recycling (Fig. 6B), and as such would inhibit its internalization in response to SNAP25Δ20 expression. However, the Cox2 (NS398) and the SphK (MPA08) inhibitors should not interfere with mPRβ internalization in response to SNAP25Δ20, although experimentally it is difficult to ascertain this. Collectively these results argue that PLA2 activity is required for mPRβ internalization and that furthermore both Cox2 and SphK activities are important to modulate oocyte maturation following mPRβ internalization.

## Discussion

mPR-dependent nongenomic signaling is an important regulator of many physiological processes in female and male reproduction, cardiovascular, neuroendocrine, neurological and immune function (3, 6, 9, 12). This raises interest in mPRs as potential therapeutic targets for hypertension and other cardiovascular diseases, reproductive disorders, neurological diseases, and cancer (12). Therefore, understanding mPR signaling is important to assess their physiological and pathological contributions. In this study, we used oocyte maturation in the frog as a well-established model for P4 nongenomic signaling.

Oocyte maturation prepares the egg for fertilization and endows it with the ability to activate and initiate embryonic development (19). As such it represents the initial step in multicellular organismal development that precedes fertilization. It is therefore not surprising that oocyte maturation involves multiple signaling cascades to mediate both the completion of meiosis -the so-called nuclear maturation- to generate a haploid gamete, and the extensive cellular differentiation of the oocyte that is needed to support development, and includes protein synthesis, membrane remodeling, and Ca^2+^ signaling differentiation that supports the block of polyspermy and resumption of meiosis at fertilization (19).

In *Xenopus* oocytes, the release of the long-term meiotic arrest can be triggered by P4. In fact, *Xenopus* oocyte maturation is one of the most studied examples of nongenomic P4 signaling and represents a well characterized system to define the signaling cascade downstream of P4. It is well established that P4 in the oocyte activates two parallel and interdependent kinase cascades that ultimately culminate in MPF activation (20). Despite many studies over the past decades, the earliest signaling steps downstream of mPR have remained elusive. An interesting recent development shows that clathrin-dependent endocytosis of mPRβ is necessary and sufficient for its signaling and the induction of oocyte maturation even in the absence of P4 (14). It therefore appears that the signaling endosome acts as a hub to activate the multiple pathways required to release meiotic arrest, resume meiosis, and prepare the egg for fertilization.

An untested assumption is that P4 induces oocyte maturation primarily through mPRβ since its knockdown or anti-mPRβ antibodies block maturation (14, 17). However, the oocyte expresses other P4 receptors including ABHD2 that may contribute to P4 signaling. Here we show that knockdown of ABHD2 blocks oocyte maturation and that this is rescued effectively by ABHD2 overexpression but not by mPRβ overexpression. ABHD2 contains a lipase and an acyltransferase motif. The lipase domain is required for oocyte maturation but not the acyltransferase domain. Based on this finding, we undertook an unbiased lipidomics approach to better define metabolic alterations driven by ABHD2 lipase activity. P4 resulted in a broad decrease in glycerophospholipid and sphingolipid species, with the enrichment of few lipid messenger end products, namely prostaglandins, LPA and S1P (Fig. 6J). This is intriguing as all these lipid messengers act through GPCRs. We in fact validate S1P action through the S1P receptor, which is a GPCR, to support oocyte maturation. These findings argue that P4 activates ABHD2 lipase activity to generate lipid messengers that in turn stimulate various GPCRs that regulate the multiple aspects of oocyte maturation. This would nicely explain the standing controversy in the field as to whether mPRs are GPCRs or not. Should the findings from the oocyte hold in other systems, it would argue that P4 activates lipid catabolism to generate lipid messenger that then act through GPCRs. Thus, nongenomic P4 signaling would involve both non-GPCR and GPCR signaling modalities in series.

An intriguing finding of the current study is that the PLA2 activity of ABHD2 is activated through its interaction with mPRβ (Fig. 6J). Although ABHD2 and mPRβ interact at rest, this interaction is enhanced following P4 treatment. This is the first report showing that ABHD2 possesses PLA2 activity and that this PLA2 activity requires interaction with mPRβ. The PLA2 family is large and complex with over 50 members that have been classified based on enzymatic activity, co-factor dependence, subcellular distribution, and structure (37, 46); and include the secreted, cytosolic, Ca^2+^-independent, lysosomal, platelet-activating factor acyl hydrolase, and some members of the α/β hydrolase domain (ABHD) family (47).

ABHD2 has been shown to act as a 2-AG lipase to activate sperm (29, 39). Interestingly, we find no evidence for such an activity in response to P4 based on our lipidomics analyses in the oocyte. This raises the interesting possibility that ABHD2 lipase specificity may be regulated by protein-protein interactions in a cell specific fashion, but this remains to be tested.

In addition to the generation of lipid messengers that branch out P4 signaling at its onset into multiple signaling cascades through GPCRs, the ABHD2 PLA2 activity hydrolyzes phospholipids to generate AA that is then metabolized to PGs through the activity of Cox2. The hydrolysis of membrane phospholipids generates lysophospholipids with a single acyl chain that tend to have a conical shape. Such conical phospholipids are prone to inducing membrane curvature (48) as the latter is determined by the size of the lipid headgroups and hydrophobic tails (49). Conical lipids have been shown to play important roles in membrane fusion events both *in vitro* and *in vivo* (50–52). It is therefore tempting to postulate that enrichment in lysophospholipids following ABHD2- PLA2 activation generates spontaneous membrane curvature, which can be sensed by the clathrin endocytic machinery to induce mPRβ endocytosis and initiate signaling and maturation (Fig. 6J).

Collectively our findings define the earliest steps in nongenomic P4 signaling to release the oocyte meiotic arrest and prepare the egg for fertilization (Fig. 6J). Furthermore, they show for the first time that two heterologous receptors previously implicated in nongenomic signaling, mPRβ and ABHD2, function as coreceptors to induce PLA2 activity that is required for both endocytosis and further downstream signaling. Nongenomic P4 signaling is widespread and mediates reproductive, neurological, and other functions. Therefore, these findings are likely to have broad implications throughout biology.

## Materials and methods

### Reagents and Primers

A list of the antibodies, siRNA, chemicals and other reagents used is provided in Supplemental Table 7. Primers and sense/antisense oligos used are listed in Supplemental Table 8. Details of the different clones used is provided in Supplemental Data.

### Xenopus laevis oocytes

All animal procedures and protocols were performed in accordance with the University of Weill Cornell Medicine-Qatar Institutional Animal Care and Use Committee. Stage VI *Xenopus laevis* oocytes were obtained as previously described (53). Oocytes were maintained in L-15 medium solution (Sigma-Aldrich Cat# L4386) supplemented with HEPES (Sigma-Aldrich Cat# H4034), 0.1% (v/v) of penicillin/streptomycin stock solution (ThermoFisher Scientific Cat# 15140-122) and 0.1% (v/v) of gentamycin (EMD Millipore Cat# 345814-1GM) at pH 7.6. The oocytes were used 24 to 72 hours after harvesting and injected with RNAs or sense/antisense oligos and kept at 18°C for 1-2 days to allow for protein expression or knockdown. After treatment with progesterone at 10^-7^ M, 10^-5^ M or 10^-4^ M (as indicated), GVBD was detected visually by the appearance of a white spot at the animal pole. For the inhibitor’s studies, oocytes were preincubated for 2 hours with different concentration of the inhibitors, followed by overnight incubation with P4 at 10^-7^ M or 10^-5^ M concentration. Co-immunoprecipitation, Western blot, and oocyte imaging are described in supplemental data.

### Metabolomic analyses

Untargeted metabolomics were performed on single oocytes or a group of 10 oocytes per condition as described in Supplemental Data. For measurements of metabolites in a group of 10 oocytes, 3 experiments using 3 independent female frogs were performed. Each experiment was conducted in 5 sample replicates per condition per frog.. Metabolites were analyzed on both the Metabolon HD4 and CLP platforms.

Tandem LC MSMS was used to measure prostaglandins and S1P from 10 oocytes using 10 replicates per condition as described in Supplemental Data. Levels of Eicosanoids, EETs and HETEs before and 30 min after P4 treatment were measured in 20 single oocytes per condition by Cayman chemicals as described in Supplemental Data.

### PLA2 assay

RNAs for mPR and ABHD2.S wt or ABHD2.S S/D/H mutants were used to generate recombinant proteins of mPR, ABHD2.S and mPR/ABHD2.S, using the rabbit reticulocytes lysates Nuclease-treated kit and the transcend^TM^ tRNA from Promega, as per the manufacturer’s instruction. These different reactions were used to measure phospholipase A2 (PLA2) activity following manufacturer’s procedures (Abcam, Cambridge, UK), in the presence of ethanol or P4 (10^-5^M). No RNA control without substrate were used as background. PLA2 activity was corrected on reticulocytes without any RNA.

We used a coupled in vitro transcription/translation kit (ALiCE^®^Cell-Free Protein Synthesis System, Sigma Aldrich) that allows the expression of difficult-to-produce proteins, such as membrane proteins, in microsomes using the pAlice02.His vector, which adds a melittin signal peptide to translocate the proteins into microsomes. To subclone mPRβ into pALICE02 vector, pSGEM-mPR was PCR amplified using the following primers: 5’ CTACCATGGCAATGACTACCGCAATCCTTGA-3’ (Forward) 5’- CTAGGTACCTCAGTGATGGTGATGGTGATGAAGTTCTTTTCTGGCCAACT-3’ (Reverse). The resulting PCR product was cut with NcoI and KpnI, gel purified, and ligated into NcoI/KpnI of pALICE02. To subclone ABHD2 in pALICE02, pSGEM-ABHD2 was PCR amplified using the following primers: 5’-ACTGATATCATGGATGCGATAGTGGAAACCC- 3’(Forward) and 5’- CTAGGTACCTCAGTGATGGTGATGGTGATGTTTATGGTCAGACTCAGCAGCC-3’(Reverse). The resulting PCR product was cut with EcoRV and KpnI, gel purified, and ligated into pALICE02 cut with NcoI and Klenow treated to produce a blunt end then cut with KpnI. All constructs were verified by DNA sequencing and by analytical endonuclease restriction enzyme digestion. The pAlice02 vector allows for the expression of C-terminally His-tagged mPRβ and ABHD2. Briefly, DNA was added to the pALiCE lysates, followed by incubation at 25°C for 48 hours with constant shaking at 700 rpm. The expression of ABHD2 and mPRβ were confirmed by western blot, before proceeding to the PLA2 assay.

### Statistics

Data are presented as mean ± SEM. Each set of experiments was at least repeated 3 times. Groups were compared using the Prism 9 software (GraphPad) using the statistical tests indicated in the figure legend. Statistical significance is indicated by p values (ns, not significant; *** p ≤ 0.001; ** p ≤ 0.01; * p ≤ 0.05).

## Acknowledgments

We are grateful to the Bioinformatics and Virtual Metabolomics Core at Weill Cornell Medicine Qatar for their support in the metabolomics and lipidomic studies. We also thank the Vivarium and Microscopy Cores at WCMQ for their support in multiple experiments. This work as well as the Cores are supported by the Biomedical Research Program at Weill Cornell Medical College in Qatar (BMRP), a program funded by Qatar Foundation. Additional funding was provided by NPRP-Standard (NPRP-S) 13th Cycle grant 13S-0206-200274 from the Qatar National Research Fund (a member of Qatar Foundation). The findings herein reflect the work and are solely the responsibility of the authors. This work was also supported by NIH R01AR076029 and NIH R21ES032347 to QC.

## Supplemental Figure Legends

**Supplemental Figure 1.**
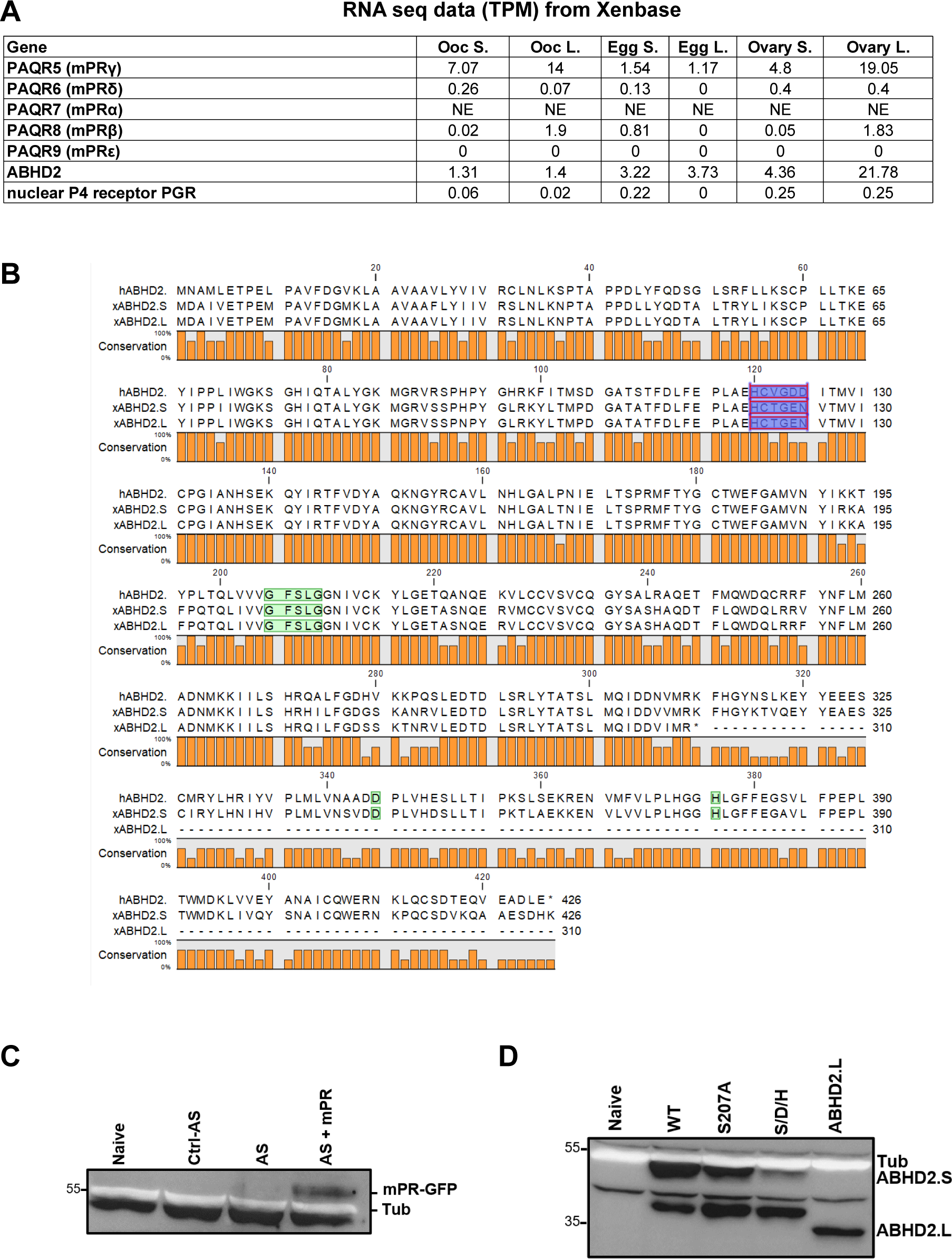
**A.** Table representing the number of transcripts per million (TPM) of the different progesterone receptors in *Xenopus laevis* oocytes (Ooc. S., and Ooc. L.), eggs (Egg S., and Egg L.) and the ovary. Being tetraploid, *Xenopus laevis* expresses two different versions of each gene; S and L. the data are retrieved from Xenbase. **B.** Alignment of *Xenopus laevis* ABHD2.L and ABHD2.S amino acids sequences with human ABHD2. Residues implicated in the lipase activity of ABHD2 are highlighted in green, whereas the acetyltransferase domain is highlighted in purple. **C.** Representative WB of mPR-GFP proteins expression in naïve, control antisense (Ctrl AS), ABHD2 antisense (AS), and AS+mPR-GFP injected oocytes. Tubulin is shown as a loading control. **D.** Representative WB of ABHD2.S protein expression levels in naïve oocytes and in oocytes injected with mRNA to overexpress ABHD2 wildtype (WT), ABHD2.S S207A, ABHD2.S S207A/D345A/H376A (S/D/H). Tubulin is shown as a loading control.

**Supplemental Figure 2.**
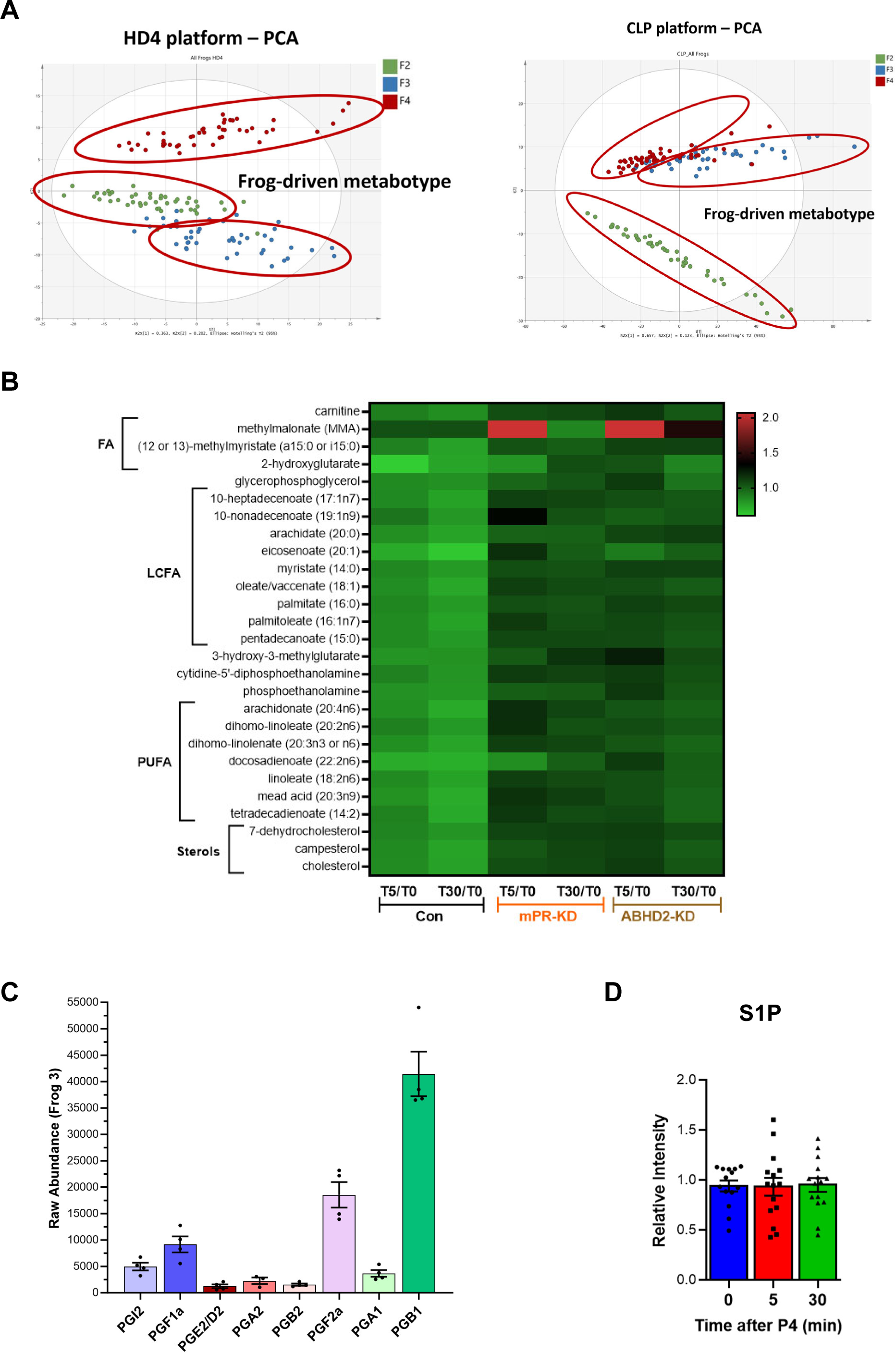
**A.** Frog-driven metabotype as shown from the HD4 (*left panel*) and CLP platforms (*right panel*). **B.** Data generated from the HD4 platform illustrated by the heatmap data of fold changes for individual metabolites that were changed significantly (p<0.05) at either the 5 min (T5) or 30 min (T30) time points in response to P4 as compared to untreated oocytes (T5/T0 and T30/T0) for naive (Con), mPR-KD, and ABHD2-KD oocytes. Metabolites are clustered at the levels of fatty acids (FA), long chain fatty acids (LCFA), polyunsaturated fatty acids (PUFA), and sterols. **C.** The raw abundance of individual prostaglandins species in pooled oocytes before treatment from a representative frog (#3) to illustrate relative abundance among the different PG species. **D.** S1P measurements in pooled oocytes before treatment, and at 5 and 30 min after progesterone addition. The data are shown as relative intensity normalized to the average of before treatment levels.

**Supplemental Figure 3.**
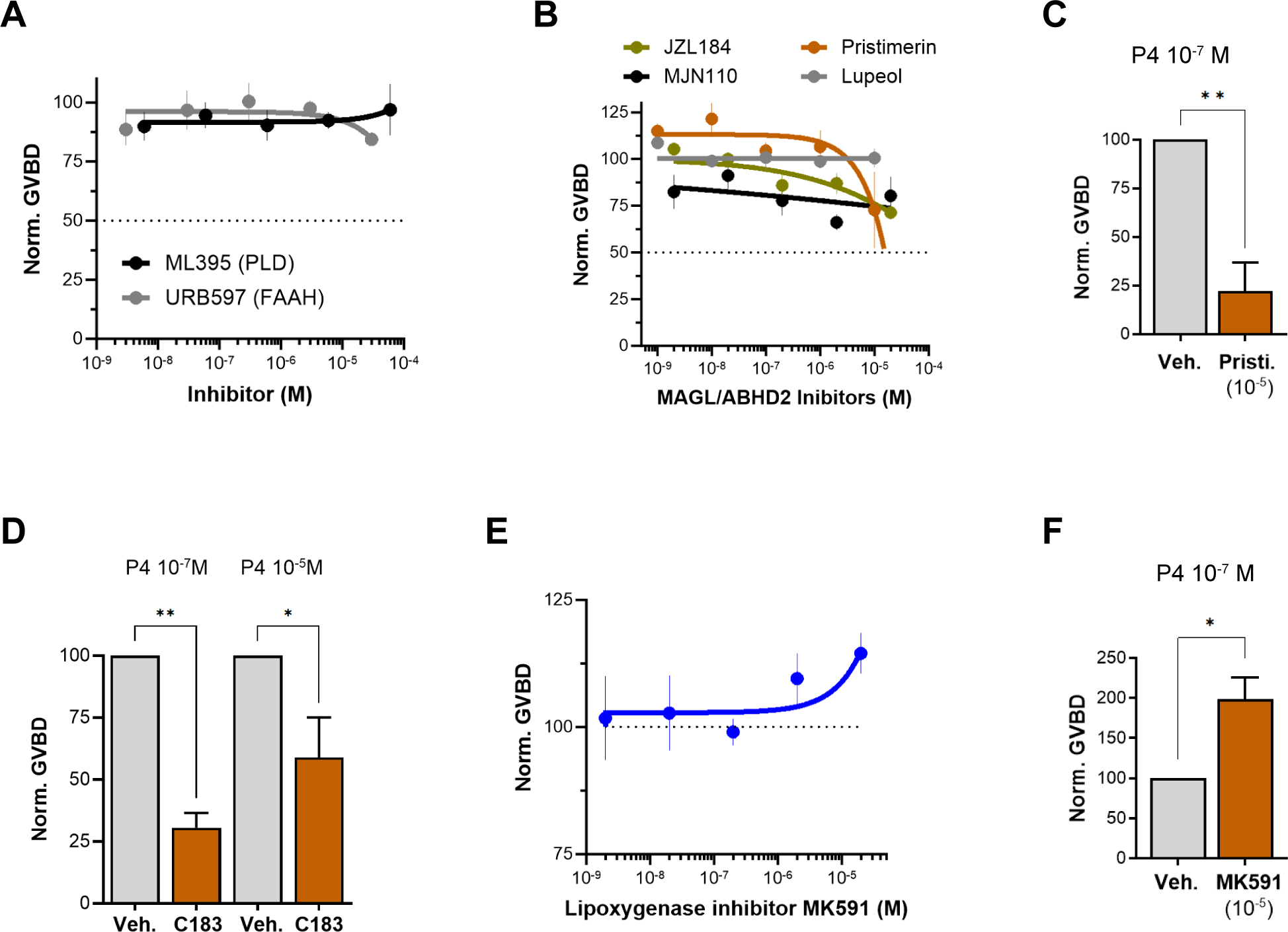
**A/B/E. Oocyte maturation inhibition dose response**. Oocytes were pre-treated for 2 hours with vehicle or with increasing concentrations of specified inhibitor, followed by overnight treatment with P4 at 10^-5^M. Oocyte maturation normalized to vehicle P4-treated oocytes. IC50 of each chemical compound was calculated using a nonlinear regression fit (mean ± SEM; n = 3 independent female frogs for each chemical compound experiment). **C/D/F.** Effect of pristimerin (**C**), compound 183 (10^-4^M) (**D**), and MK591 (**F**) on oocytes maturation at the indicated P4 concentration. Oocytes were pre-treated for 2 hours with the vehicle or indicated inhibitor, followed by overnight treatment with P4. Oocyte maturation was normalized to the control oocytes condition (treated with vehicle) (mean ± SEM; n = 3 independent female frogs for each chemical compound experiment, unpaired t-test).

**Supplemental Figure 4.**
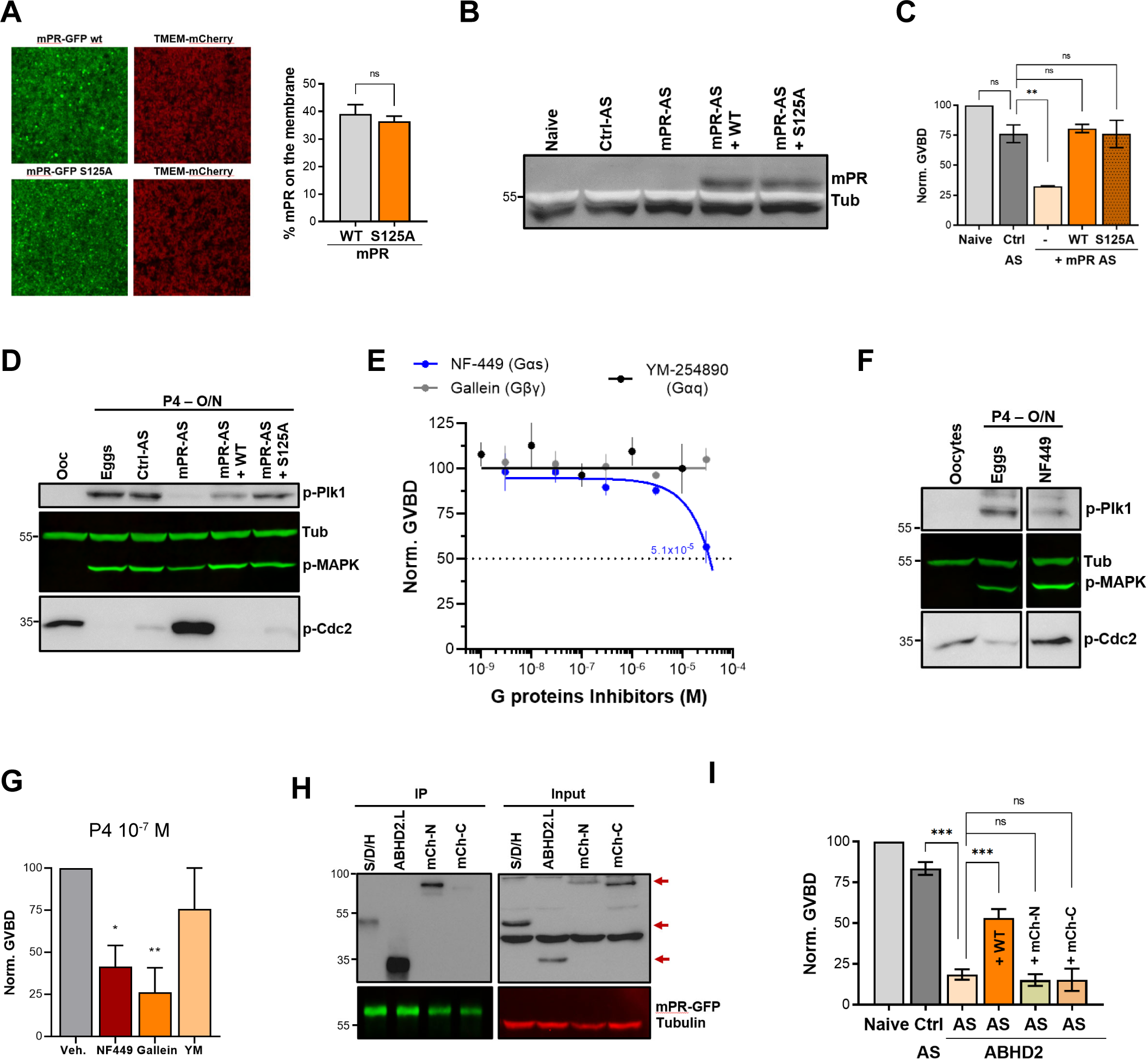
**A**. mPR S125 plasma membrane localization. Oocytes were injected with mPR-GFP wild type (WT) or the S125A mutant, along with TMEM-mCherry as a plasma membrane (PM) marker*. Left panel,* Representative confocal image of oocytes overexpression mPR-GFP WT or mPR-GFP S125A mutant along with TMEM-mCherry. *Right panel,* histogram showing the percentage of mPR-GFP WT or mPR-GFP S125A at the PM (mean ± SEM; n = 10 oocytes per clone from 2 independent female frogs, unpaired t-test). **B.** Representative WB of mPR-GFP protein expression in naïve, control antisense (Ctrl AS), mPRβ antisense (AS), and AS+mPR-GFP WT or mPR-GFP S125A mutant injected oocytes. Tubulin is shown as a loading control. **C.** Oocyte maturation in oocytes injected with control antisense (Ctrl AS) or mPRβ antisense (AS) with or without mPR-GFP WT or mPR-GFP S125A expression, and normalized to P4-treated naïve oocytes (Naive) (mean ± SEM; n = 3 independent female frogs, ordinary one-way ANOVA). **D.** Representative WB of MAPK, Plk1 and Cdc2 phosphorylation from untreated oocytes, P4 matured eggs, oocytes injected with control antisense (Ctrl AS) or mPRβ antisense (AS) with or without mPR-GFP WT or mPR-GFP S125A expression. Tubulin is shown as a loading control. **E.** Inhibitor dose response. Oocytes were pre-treated for 2 hours with vehicle or with increasing concentrations of specified inhibitor, followed by overnight treatment with P4 at 10^-5^M. Oocyte maturation normalized to vehicle P4-treated oocytes. IC50 was calculated using a nonlinear regression fit (mean ± SEM; n = 3 independent female frogs for each chemical compound experiment). **F.** Representative WB of MAPK, Plk1 and Cdc2 phosphorylation from untreated oocytes, oocytes pretreated with vehicle for 2 hours then matured overnight (O/N) by P4 at 10^-7^M (eggs), and oocytes pretreated for 2 hours with NF449 then treated O/N with P4 10^-7^M. Tubulin is shown as a loading control. **G.** Drug effect on oocyte maturation at low P4 concentration. Oocytes were pre-treated for 2 hours with vehicle or the specified inhibitor followed by overnight treatment with P4 at 10^-7^M. Oocyte maturation was normalized to the control oocytes condition (treated with vehicle) (mean ± SEM; n = 3 independent female frogs for each chemical compound experiment, unpaired t-test). **H.** Representative WB of the immunoprecipitation (IP) of mPR-GFP using lysates from oocytes over- expressing mPR-GFP with ABHD2.L, mCherry tagged ABHD2.S at its C-terminal (mCh-C) or N-terminal (mCh-N), and ABHD2.S S/D/H. The input represents expression levels of the different clones before the IP. Tubulin is shown as a loading control. The red arrows point to ABHD2.S different clones. **I.** Oocyte maturation measured in oocytes injected with control antisense (Ctrl AS) or ABHD2 antisense (AS) with or without ABHD2.S wild type (AS+WT) or the mCherry tagged ABHD2.S clones, and normalized to P4-treated naïve oocytes (Naive) (mean ± SEM; n = 3 to 5 independent female frogs, ordinary one-way ANOVA).

**Supplemental Table 8.** List of antibodies, chemicals, and reagents.

List of antibodies, chemicals, and reagents
| REAGENT | Source | Catalogue # |
| --- | --- | --- |
| **Antibodies** |  |  |
| Rabbit polyclonal anti-ABHD2 | Sigma-Aldrich | Cat# HPA005999 |
| Rabbit polyclonal anti-GFP | Living Colors | Cat# 632381 |
| Mouse monoclonal anti-SNAP25 | Biolegend | Cat# 836304 |
| Rabbit polyclonal anti-p-plk1 | Cell signaling | Cat# 5472 |
| Mouse monoclonal anti-phospho-MAPK | Cell Signaling | Cat# 9106 |
| Rabbit polyclonal anti-phospho-Cdc2 | Cell Signaling | Cat# 9111 |
| Rabbit polyclonal anti-Tubulin | Cell Signaling | Cat# 3873 |
| Rabbit polyclonal anti-Cdc25C (Ser216) | Cell signaling | Cat# 9528 |
| Rabbit polyclonal anti-S1PR3 | Bioss | Cat# BS-7541R |
| Goat anti-rabbit IgG-HRP | Jackson ImmunoResearch | Cat# 111-035-144 |
| Goat anti-rabbit IgG- IRDye® 800 | Li-COR | Cat# 925-32211 |
| Goat anti-mouse IgG- IRDye® 680 | Li-COR | Cat# 925-68070 |
| Goat anti-mouse IgG- IRDye® 800 | Li-COR | Cat# 926-32210 |
| Mouse polyclonal anti-His | Takara | Cat# 631212 |
| **Chemicals and kits** |  |  |
| Progesterone | Sigma-Aldrich | Cat# 200-350-9 |
| Pitstop® 2 | Abcam | Cat# ab120687 |
| Pitstop® 2 - negative control | Abcam | Cat# ab120688 |
| Mammalian ProteaseArrest™ | G-Biosciences | Cat# 786-331 |
| PhosphataseArrest™ | G-Biosciences | Cat# 786-451 |
| µMACS GFP Isolation Kit | Miltenyi Biotec | Cat# 130-091-125 |
| ECL™ Prime Western Blotting Detection | GE | Cat# 45-002-401 |
| The XL QuikChange mutagenesis kit | Agilent Technologies | Cat# 200516 |
| The mMessage mMachine T7 kit | ThermoFisher | Cat# AM1344 |
| Fast SYBR™ Green Master Mix | ThermoFisher | Cat# 4385616 |
| RNeasy Mini Kit | Qiagen | Cat# 74104 |
| Rabbit Reticulocyte Lysate | Promega | Cat# L4960 |
| ALiCE® | Sigma | Cat# AL0103200 |
| Trancend™ tRNA | Promega | Cat# L5061 |
| Phospholipase A2 Activity assay kit | Abcam | Cat# ab273278 |
| FAAH | Sigma-Aldrich | Cat# 341249 |
| MJN110 | Sigma-Aldrich | Cat# SML0872 |
| JZL184 | Sigma-Aldrich | Cat# J3455 |
| Pristimerin | Sigma-Aldrich | Cat# P0020 |
| Lupeol | Sigma-Aldrich | Cat# L5632 |
| ML395 | Merck | Cat# 532978 |
| cPLA2 Inh II | Sigma-Aldrich | Cat# 5305380001 |
| ACA | Sigma-Aldrich | Cat# A8486 |
| Bromo-enol lactone | Sigma-Aldrich | Cat# B1552 |
| AACOCF3 | Abcam | Cat# ab120350 |
| Mastoparan | Enzo | Cat# BML-G410-0001 |
| HA130 | Tocris | Cat# 4196 |
| MK591 | Sigma-Aldrich | Cat# 147030-01-1 |
| NS398 | Sigma-Aldrich | Cat# 349254 |
| SC-560 | Sigma-Aldrich | Cat# 565610 |
| D-erythro MAPP | Enzo | Cat# SL221 |
| L-erythro MAPP | Enzo | Cat# SL222 |
| MP-A08 | Tocris | Cat# 5803 |
| NF-449 | Sigma-Aldrich | Cat# 480420 |
| Gallein | Sigma-Aldrich | Cat# 371708 |
| YM-254890 | Cayman | Cat# 29735 |
| Org OD-02-0 | Axon Med Chem | Cat# 2085 |
| **Software** |  |  |
| Image Studio 5.2 | LI-COR Biosciences | <u>N/A</u> |
| ImageJ | Schneider et al. 2012 | <u>N/A</u> |
| Prism | GraphPad inc. | <u>N/A</u> |
| Illustrator CC | Adobe Systems Inc | <u>N/A</u> |
| Photoshop CC | Adobe Systems Inc | <u>N/A</u> |
| ZEN 2.3 black | Zeiss | <u>N/A</u> |
| Quant Studio Real-Time PCR software.Ink | ThermoFisher | <u>N/A</u> |
| Biorender | Biorender | N/A |

**Supplemental Table 9.**
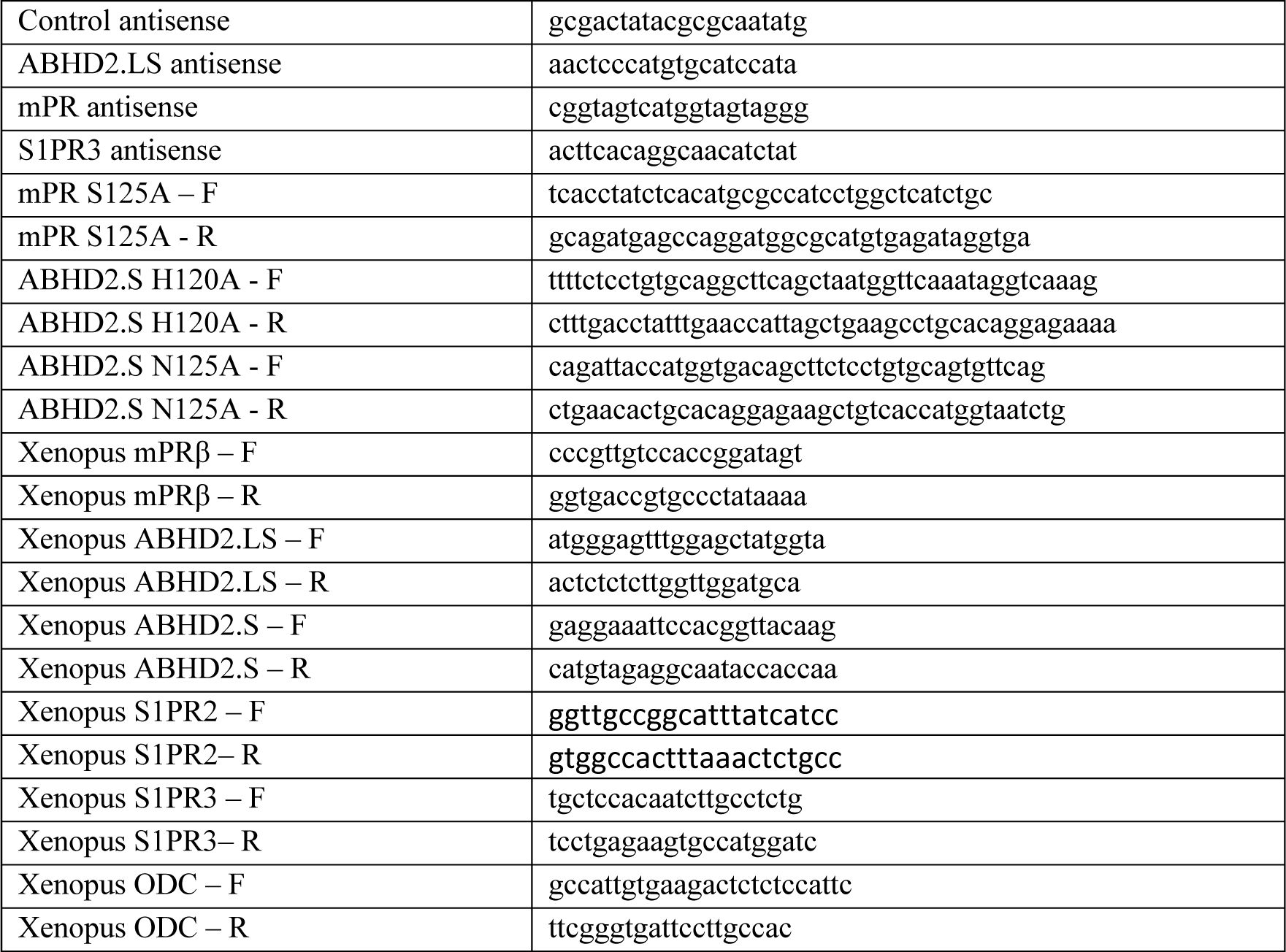
List of antisense oligonucleotides and primers.

